# Recent evolutionary origin and localized diversity hotspots of mammalian coronaviruses

**DOI:** 10.1101/2023.03.09.531875

**Authors:** Renan Maestri, Benoît Perez-Lamarque, Anna Zhukova, Hélène Morlon

## Abstract

Several coronaviruses infect humans, with three, including the SARS-CoV2, causing diseases. While coronaviruses are especially prone to induce pandemics, we know little about their evolutionary history, host-to-host transmissions, and biogeography. One of the difficulties lies in dating the origination of the family, a particularly challenging task for RNA viruses in general. Previous cophylogenetic tests of virus-host associations, including in the Coronaviridae family, have suggested a virus-host codiversification history stretching many millions of years. Here, we establish a framework for robustly testing scenarios of ancient origination and codiversification *versus* recent origination and diversification by host switches. Applied to coronaviruses and their mammalian hosts, our results support a scenario of recent origination of coronaviruses in bats and diversification by host switches, with preferential host switches within mammalian orders. Hotspots of coronavirus diversity, concentrated in East Asia and Europe, are consistent with this scenario of relatively recent origination and localized host switches. Spillovers from bats to other species are rare, but have the highest probability to be towards humans than to any other mammal species, implicating humans as the evolutionary intermediate host. The high host-switching rates within orders, as well as between humans, domesticated mammals, and non-flying wild mammals, indicates the potential for rapid additional spreading of coronaviruses across the world. Our results suggest that the evolutionary history of extant mammalian coronaviruses is recent, and that cases of long-term virus–host codiversification have been largely over-estimated.

## Introduction

Coronaviruses are RNA-viruses of the family Coronaviridae, comprising positive-sense and single-stranded viruses that have the largest genomes among nidoviruses (1, 2). As with several other RNA viruses, they may cause diseases in humans and other animals (3). Depending on the taxonomic arrangement, seven (4–6) or eight (7) species of coronaviruses infect humans, three of which being pathogenic: the SARS-CoV (8, 9), the MERS-CoV (10), and the SARS-CoV-2 (11). The latter is at the origin of the recent COVID-19 pandemic that infected more than 775 million people and caused the death of more than seven million (12). Coronaviruses’ high frequency of recombination (13), broad host range, and high mutation rates (7) make them especially prone to causing yet future diseases. Nevertheless, their evolutionary history and biogeography are very poorly understood. Resolving the evolutionary origins of Coronaviridae, understanding how they diversified, and characterizing their geographic diversity patterns would facilitate attempts to predict future zoonoses (7, 14, 15).

Coronaviruses infect mammals, birds, and fishes (2), although they predominate in mammalian species (16–20). A consensus exists on the taxonomic segregation of four genera within Coronaviridae: Orthocoronavirinae, namely Alpha-, Beta-, Gamma- and Deltacoronavirus (2, 21). Alpha- and betacoronaviruses are found exclusively in mammals, while delta- and gammacoronaviruses infect mostly birds but also mammals to a lesser extent (20, 22, 23). Coronaviruses are most numerous and genetically diversified in mammals (2, 23), in particular bats, suggesting a mammalian origin in bats (2, 20, 23, 24), although this remains to be tested.

The timing of origination of the Coronaviridae family is debated, with results that vary by several orders of magnitude. Woo et al (23) found a recent origin, around 10 thousand years ago. This dating was obtained by sequencing the well-conserved RNA-dependent RNA-polymerase (RdRp) genome region of representatives of all four coronavirus genera, and fitting to these sequences a neutral nucleotide-based substitution model with an uncorrelated log-normal relaxed clock (25) calibrated with serial samples. This calibration provided a mean substitution rate estimate of 1.3 x 10-4 substitutions per site per year. Wertheim et al. (26) used this estimate and the same genome region (RdRp), but with a codon-based substitution model accounting for the effect of selection. Indeed, purifying selection can lead to an underestimation of viral origins when not accounted for (26, 27). They found an ancient origin, around 293 (95% confidence interval, 190 to 489) million years ago (26). More recently, Hayman & Knox (28) obtained similar results, but using the splitting times of hosts as constraints, therefore assuming a priori that coronaviruses codiversified with their hosts.

More generally, dating the phylogenies of RNA virus families is a difficult task (29). While for some of them dated calibration points can be used, based on orthologous copies of endogenous virus elements (EVEs) present in the genomes of related mammalian species with known times of divergence (30), in many others, including in the Coronaviridae, such elements have not been found (31). Despite the difficulty in dating viral families, it has been proposed, from cophylogenetic analyses investigating the congruence of the host and viral phylogenetic trees (32), that vertebrate-associated RNA viruses have codiversified with their hosts over hundreds of millions of years (31, 33; but see 34; see Box S1 for a clarification of out terminology). Indeed, RNA virus phylogenies tend to mirror that of their hosts; for example, closely-related coronaviruses infect closely-related mammals (e.g. (28)). However, a major caveat is that such cophylogenetic signals can emerge when viruses diversify by host switches preferentially occurring among closely-related hosts, in the absence of any cospeciation event (35, 36).

Event-based cophylogenetic methods can in principle identify cospeciation and host switches events (32, 35), but their behavior in the presence of diversification by preferential host switches is not well understood. Under a perfect codiversification scenario, host and symbiont phylogenies would be identical. Events of host switches, duplications and losses induce mismatches, and cophylogenetic methods aim to identify parsimonious sets of events that allow “reconciling” the two phylogenies (35, 37). However, most of these methods rely entirely on tree topology (and not branching times), such that time-inconsistent host switches between non-contemporary host lineages are allowed during the reconciliation. In the presence of preferential host switches, these methods may thus favor biologically unrealistic reconciliations that involve cospeciation events and ‘back-in-time’ host switches to reconciliations that involve more frequent contemporary host switches. This would have remained unnoticed, unless users of the methods specifically looked at the time consistency of the inferred host switches, which is usually not done.

Here, we establish a framework for testing scenarios of ancient origination and codiversification versus recent origination and diversification by host switches that combines probabilistic cophylogenetic models and biogeographic analyses (Fig 1). We then apply this framework to the Coronaviridae-mammals association. We assemble a dataset of all mammalian hosts of coronaviruses and a complete association matrix between host species and species-like Operational Taxonomic Units (sOTUs) of coronaviruses, as well as geographic repartition of Coronaviridae and their mammalian hosts. We construct a new Coronaviridae tree based on a recent proposition for the use of a well-conserved region of their RNA genome (38, 39). Under the ancient origination scenario (Fig 1A), long-term vertical transmission of Coronaviridae within mammalian lineages could lead to events of mammal-coronavirus cospeciations. Coronaviruses’ diversification would then be modulated by both cospeciations and horizontal host switches from one mammalian lineage to another (26, 31). The most recent common ancestor of coronaviruses could even have infected the most recent common ancestor of mammals and birds (26). Under the recent origination scenario (Fig 1B), codiversification with hosts is virtually impossible, and coronaviruses’ diversification would then be largely dominated by recent host switches. Expectations for the output of reconciliation and biogeographic analyses under these different scenarios, as well as a scenario of random associations, are explained in Fig 1. We identify the likely origination of coronaviruses in the mammalian tree, quantify the frequency of cospeciation and host-switching events, and locate these host switches, therefore identifying ‘reservoirs’ of Coronaviridae and potential transmission routes across mammals.

**Figure 1.**
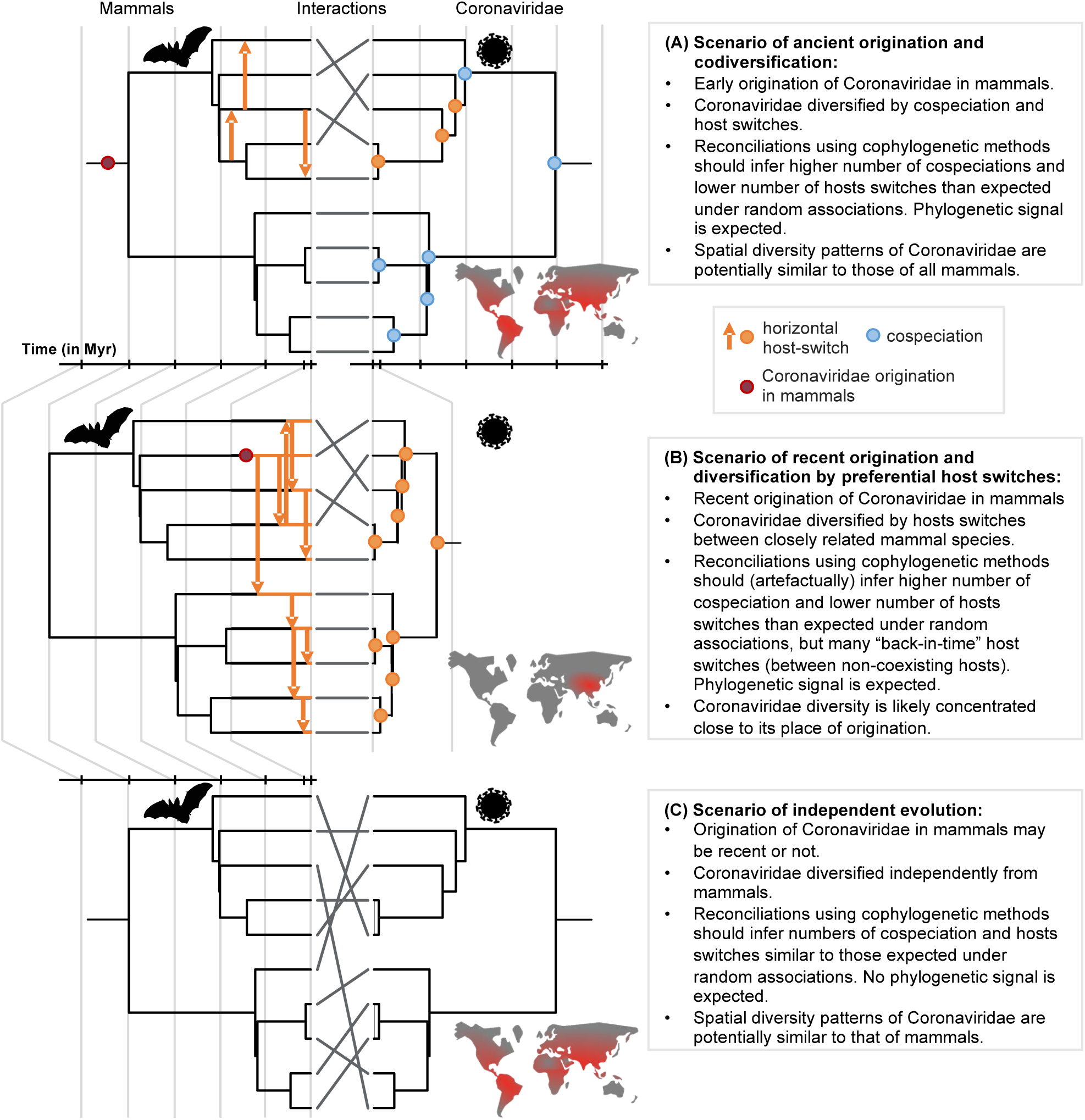
A framework for testing scenarios of virus-host evolution, illustrated with the example of Coronaviridae and their mammalian hosts: In (A), a scenario of ancient origination and codiversification; in (B) a scenario of recent origination and diversification by preferential host switches; and in (C) a scenario of independent evolution. For each scenario, we indicate the associated predictions in the grey boxes. Contrary to scenario C, both scenarios A and B are expected to generate a cophylogenetic signal, *i.e.* closely-related coronaviruses tend to infect closely-related mammals, resulting in significant reconciliations when using topology-based probabilistic cophylogenetic methods, such as the undated version of ALE, Jane, or eMPRess. However, we expect scenario B to be distinguishable from scenario A in terms of the time consistency of host-switching events. Under scenario B, cophylogenetic methods wrongly estimate a combination of cospeciations and “back-in-time” host switches (see Methods & Results). We also expect different biogeographic patterns under the different scenarios, as illustrated by the maps, where the color gradient represents diversity levels (red: high diversity, grey: low diversity).

## Results

### Mammal-coronavirus associations

By screening the 46 sOTUs of Coronaviridae identified by Edgar et al. (38) in public databases, we found 35 that were associated with mammalian hosts. Our trees of these 35 sOTUs support a well-defined split between alphacoronaviruses and the other genera, regardless of the phylogenetic method used (Fig 2; *SI Appendix*, Fig. S1). Overall, alphacoronaviruses form a monophyletic clade, delta- and gammacoronoviruses form sister clades, with the main uncertainty being on the placement of their ancestor in relation to betacoronaviruses (i.e. as a sister to a monophyletic Beta-clade (Fig. 2) or within the Beta-clade (*SI Appendix*, Fig. S1).

**Figure 2.**
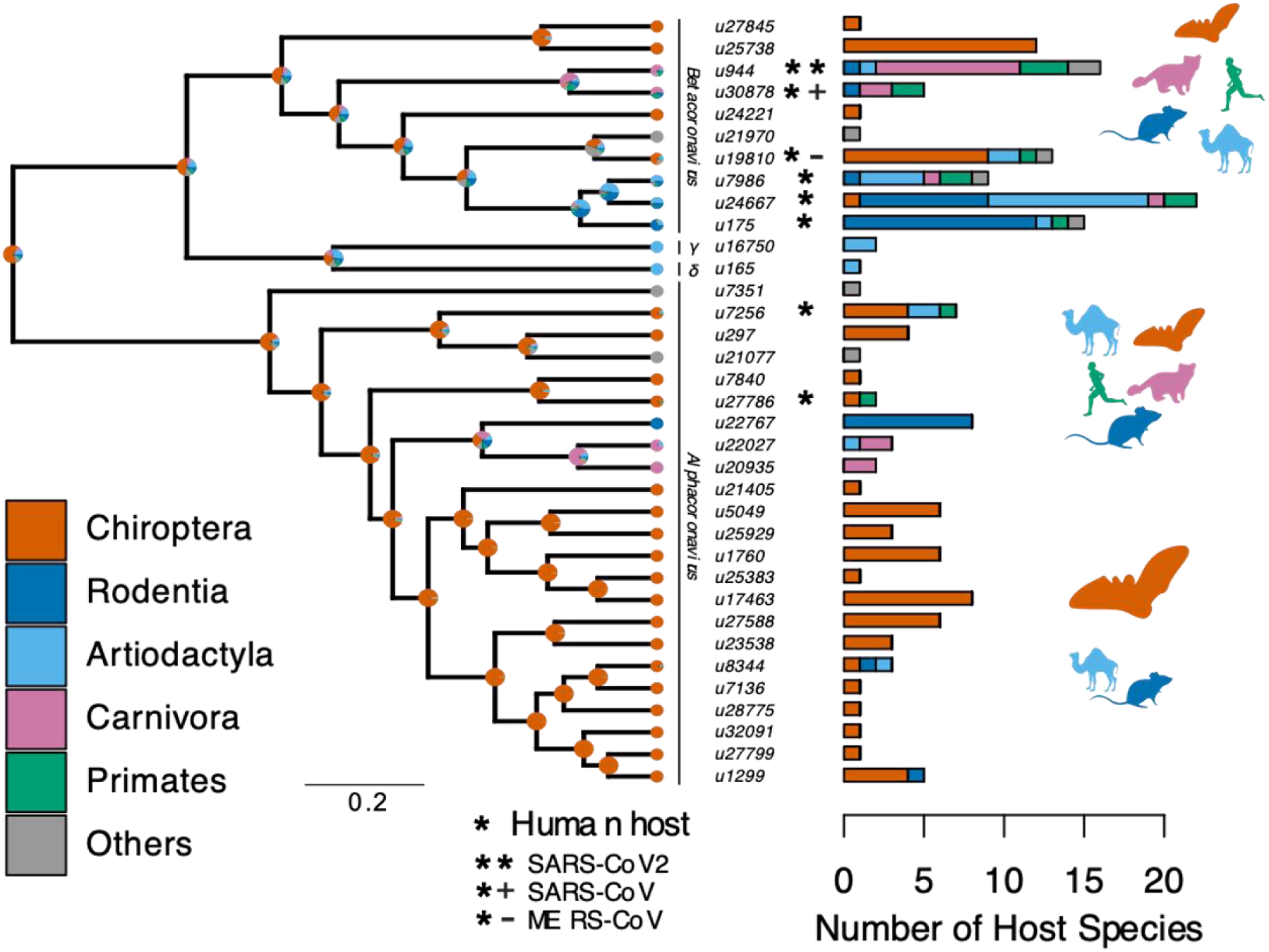
Species-level relationships among coronaviruses and their associated mammalian hosts. The Maximum Clade Credibility phylogenetic tree of coronaviruses, reconstructed with BEAST2 based on 150-aa palmprint amino acid sequences of the RdRp gene, is shown on the left. sOTUs of Coronaviridae followed the definition of the Serratus project. The branching order of four genera of coronaviruses, Beta, Gamma, Delta, and Alphacoronaviruses, is shown. Bar scale is in units of aa substitution. On the right, a barplot gives the number of total mammalian host species and the number of host species by main mammalian order. Ancestral states on the left were obtained for illustrative purposes with the make.simmap function of the phytools R package (Revell 2012). Mammal silhouettes taken from open-to-use sources in phylopic.org, detailed credits given in SI Appendix Table S6.

We found that mammalian hosts of coronaviruses belong to 31 families and 10 orders of mammals, and are widely distributed throughout the mammalian phylogeny (Fig. 2, SI Appendix, Fig. S2). Most mammalian hosts are bats (Chiroptera - 55 species), followed by rodents (Rodentia - 22 species), artiodactyls (Artiodactyla - 15 species), carnivores (Carnivora - 11 species), and primates (Primates - 5 species). Five other orders have at least one representative species: Eulipotyphla (4), Lagomorpha (1), Perissodactyla (1), Pholidota (1), and Sirenia (1). The number of mammalian hosts per coronavirus’ sOTU varies across the Coronaviridae tree, ranging from 1 to 22 species, with an average of 4.94 (Fig. 2). Of the 35 sOTUs, 23 are found in at least one bat species and 17, mostly in alphacoronaviruses, are found exclusively in bats (Fig. 2). Eight sOTUs are found in humans, six of which, including the three pathogenetic sOTUs, are betacoronaviruses. Betacoronaviruses infect a larger average number of hosts and a larger diversity of non-bat species than alphacoronaviruses. Twenty-two coronaviruses occur in more than one species; of those, 11 are found in multiple orders (Fig. 2; *SI Appendix*, Fig. S3) and 11 in multiple species of a single order (*SI Appendix*, Fig. S3). A simple visualization of the mammal-coronavirus interaction network indicates that mammals from the same orders tend to be infected by the same coronavirus’ sOTUs, that bats tend to host a specific set of coronaviruses, and the centrality of humans in the network (Fig. 3).

**Figure 3.**
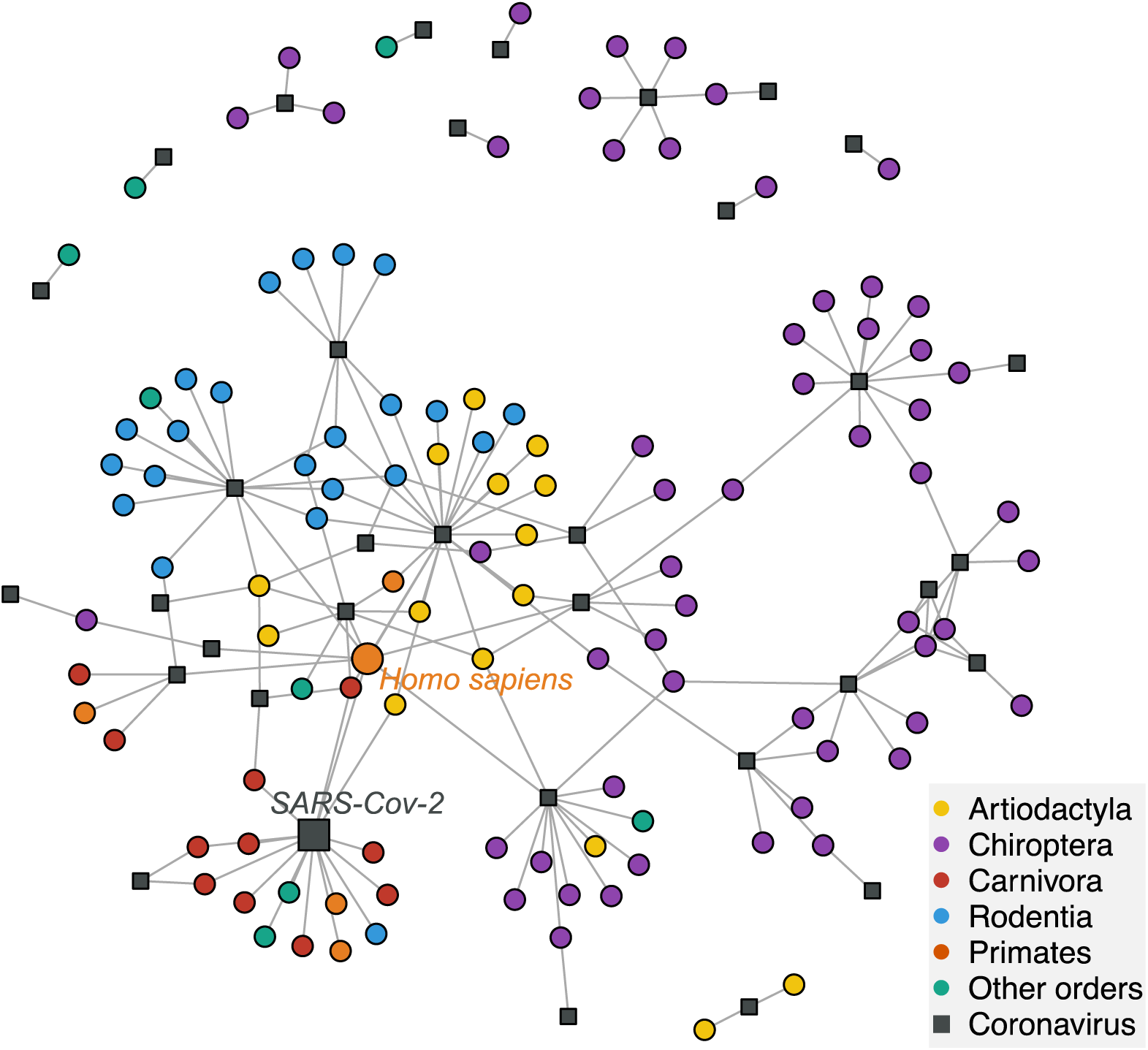
A network visualization of mammal-coronavirus interactions reveals the presence of phylogenetic signal, the isolation of bats, and the centrality of humans. Species-level network representation of the interactions between mammal species and coronavirus sOTUs. Colored round nodes represent mammal species (colors indicate the mammalian order) and grey squared nodes correspond to coronavirus sOTUs. The position of the nodes reflects their similarity in interaction partners, i.e. the tendency of clustering of mammals belonging to the same order can be interpreted as the presence of phylogenetic signal in species interactions. Humans and SARS-Cov-2 are presented using bigger nodes. The plot was obtained using the Fruchterman-Reingold layout algorithm from the igraph R-package.

### Phylogenetic signal in coronavirus infections

We first tested whether closely-related coronaviruses tend to infect closely-related mammals. A negative answer to this question would suggest that the diversification of Coronaviridae is independent of mammalian history, excluding the scenarios of codiversification or diversification per preferential host switches (Fig. 1). To the contrary, we found a significant phylogenetic signal for the overall association between coronaviruses and mammals (Mantel test: r= 0.38; *P*= 0.0001) and vice-versa (r= 0.29; *P*= 0.0001), after accounting for the confounding phylogenetic signal in the number of partners (40). Mantel tests across sub-clades of both phylogenies revealed that this overall phylogenetic signal is linked to phylogenetic signal in the deep nodes of the Coronaviridae and mammal phylogenies rather than at shallow phylogenetic scales (*SI Appendix*, Fig. S4), consistent with the order-level pattern observed in the mammal-coronavirus interaction network (Fig. 3). This pattern could arise from ancient codiversification followed by un-preferential host switches, or from recent host switches preferentially occurring between hosts from the same high-level taxonomic grouping (such as mammalian orders). We also found that closely related coronaviruses tend to infect a similar number of hosts (r= 0.29; *P*=0.002), while closely related mammals do not tend to host a similar number of distinct coronaviruses (r= 0.04, *P*=0.1), suggesting that coronaviruses’ specificity towards hosts is evolutionarily conserved while hosts’ specificity to coronaviruses is not.

### Diversification dynamics of coronaviruses

To further investigate the hypotheses of ancient codiversification versus recent host switches, we used a probabilistic cophylogenetic model, the amalgamated likelihood estimation (ALE – (41)), that reconciles the host and symbiont phylogenies using events of cospeciations, host switches, duplications, or losses, while accounting for phylogenetic uncertainty in the symbiont phylogenies and undersampling of the host species (37, 41, 42). The main version of ALE we used is an “undated” version that accounts for topology but not branch lengths, as the dated version did not perform well on our data (see Methods). Time-inconsistent host switches are thus allowed during the reconciliation. If the scenario of ancient diversification holds, we expect to find reconciliations requiring more cospeciations and fewer host switches than expected under a scenario of independent evolution (hereafter referred to as ‘significant reconciliation’), and few time-inconsistent switches (Fig. 1A). Under the alternative scenario of recent origination and diversification by preferential host switches, we also expect to infer a significant reconciliation, but with many time-inconsistent switches, as the algorithm tends to explain the cophylogenetic signal in the interactions by cospeciation events (Fig. 1B). We indeed found a significant reconciliation between the Coronaviridae and the mammalian trees, confirming the non-independence of their evolution, which we evaluated by randomly shuffling mammal species across the full tree or within biogeographic regions (*SI Appendix*, Fig. S7). ALE reconciliations inferred average numbers of 145 cospeciations, 65 losses, 0 duplication, and 92 host switches. Without investigating the time-consistency of the host switches, we would conclude that there are almost 1.5 more diversification events of Coronaviridae that are related to ancient codiversification rather than host switches. However, on average 20% of the inferred host switches are time-inconsistent, including “back-in-time” host switches of >50 Myr (*SI Appendix*, Fig. S8). Similar results were observed when accounting for the uncertainty in the branching time estimates (*SI Appendix*, Fig. S8), which suggests instead that extant Coronaviridae originated recently and diversified by frequent preferential host switches.

The majority (64%) of the reconciliations found an origination of coronaviruses within bats, in particular within the Pteropodidae family (Fig. 4A-C). By comparison, only 19% found an origination in rodents, 6% in artiodactyls, and 2% in carnivores. In addition, we no longer found an origination in bats when randomly shuffling the dataset (*SI Appendix*, Fig. S5,S6), suggesting that this result is not artifactual. We checked the interpretation of our results by simulating the two scenarios of (i) ancient origination in the ancestors of bats followed by codiversification and (ii) recent origination in an extant bat species and a subsequent diversification by preferential host switches. On the first set of simulations, ALE correctly inferred an origination in bats and very few time-inconsistent switches (2% +/- 2%), which seems to be the basal expected proportion of time-inconsistent switches under a scenario of codiversification (*SI Appendix*, Fig. S12). On the second set, ALE correctly inferred an origination in bats, although with lower confidence, and a large fraction (∼20%) of time-inconsistent host switches, similar to what we observed for Coronaviridae. These results therefore indicate a scenario of recent origination of coronaviruses in bats followed by diversification by preferential host switches.

**Figure 4.**
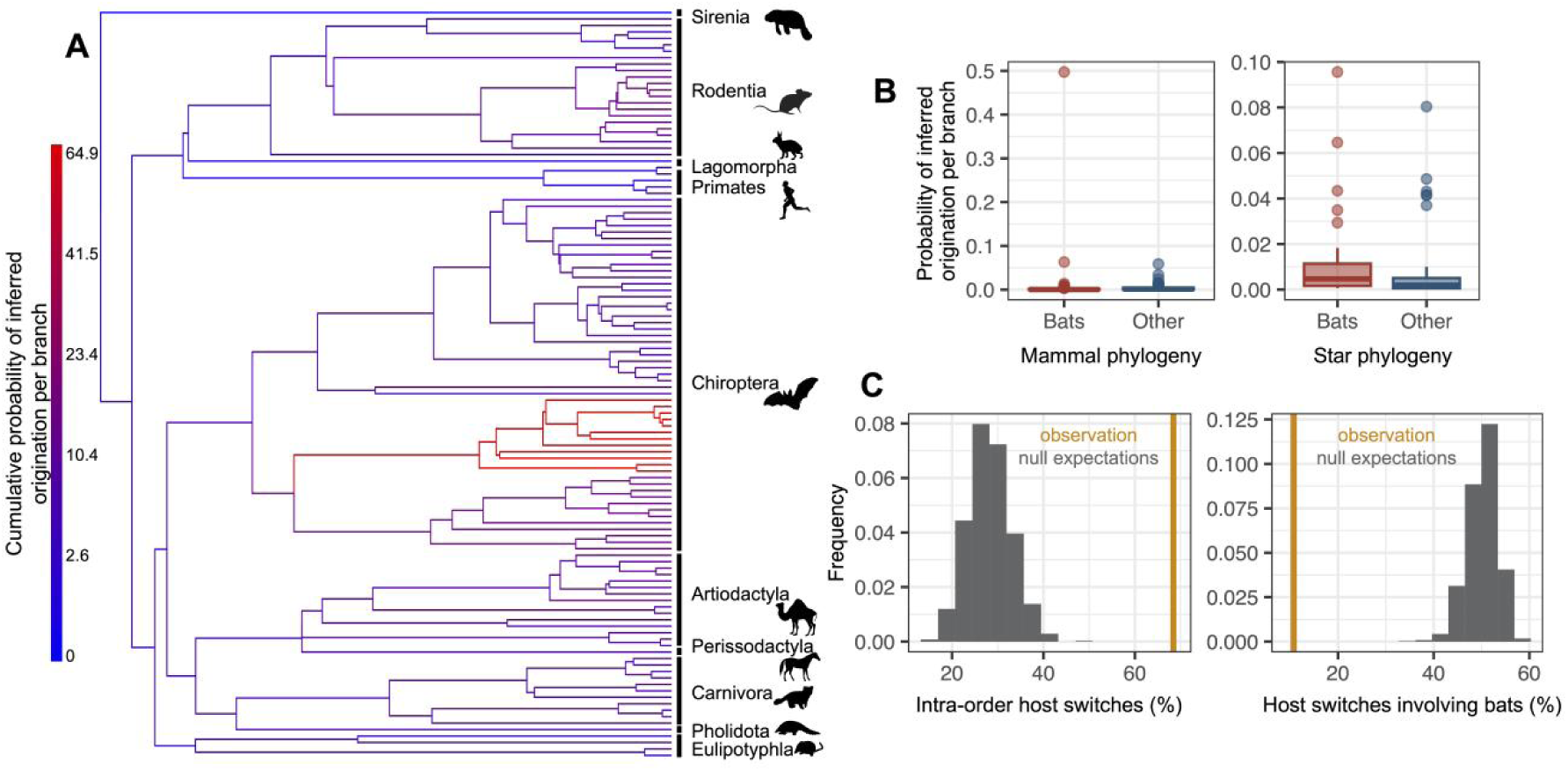
The origination of coronaviruses in mammals is estimated among bats, which tend to form a closed reservoir. (A) Phylogenetic tree of the mammals with branches colored as the percentage of ALE reconciliations which inferred this branch or its ancestral lineages as the origination of coronaviruses in mammals. Red branches are likely originations, whereas blue branches are unlikely. (B) Boxplots recapitulating the probability of inferred origination per branch in bats *versus* other mammal orders, with ALE applied on the original mammal tree (left panel) or on the mammal tree transformed into a star phylogeny (right panel), therefore assuming an origination in extant species. (C) Distributions of the percentages of host switches occurring within mammalian orders (left panel) and between-orders involving bats (right panel). Observed values (in orange) are compared to null expectations if host switches were happening at random (in grey). Mammal silhouettes taken from open-to-use sources in phylopic.org, detailed credits given in SI Appendix Table S6.

To investigate this scenario in more detail, we gradually applied a tree transformation to the mammalian phylogeny, which excludes the possibility of an ancient origination happening earlier than a given time. We found that we had to impose a very recent time of origination (younger than 5 Myr) to obtain few time-inconsistent switches (*SI Appendix*, Table S1). We thus carried out our follow-up analyses with a mammals’ tree transformation (star phylogeny) that assumes an origination in an extant mammalian lineage, such that coronavirus diversification is explained entirely by host switches between extant mammalian species. Simulations validated this approach in terms of properly inferring originations and identifying preferential host switches (*SI Appendix*, Fig. S13). Applied to the data, the approach inferred a high probability of origination in bats (56%, Fig, 4B-C, *SI Appendix*, Fig. S6), far more likely than in other mammalian orders (artiodactyls: 18%, rodents: 7%, carnivores: 5%). Simulations confirmed that our results did not spuriously arise because of higher coronaviruses diversification in bats compared to other mammalian orders (Fig. S14).

Following the origination in bats, our approach based on the star phylogeny inferred a scenario of diversification by preferential host switches: 68% of the inferred host switches happened within mammal orders (Fig 4D, *SI Appendix*, Fig. S10), whereas we would expect on average only 28% of within-order host switches if happening at random. We also inferred more-than-expected host switches between closely related mammal orders (e.g. between Artiodactyla and Perissodactyla) and between the order containing humans (Primates) and those of their domesticated animals, such as Artiodactyla and Carnivora (*SI Appendix*, Fig. S10, Table S2). In contrast, host switches were five times less numerous than expected by chance between bats and other orders (10.7%, against 50.2% on average if host switches were randomly distributed, Fig. 4D), in particular Artiodactyla and Rodentia (*SI Appendix*, Fig. S11, Table S2). When occurring, host switches from bats often occurred toward humans (1.9 host switches per reconciliation on average) or toward urban-living and/or domesticated animals, such as rats, camels, or pigs (>1 host switch on average; *SI Appendix*, Table S3). Host switches to humans occurred mostly from domesticated mammals (camels, pigs, dogs), the house shrew and the house mouse, then followed by Asian palm civets, and lastly by bats and other rodents (*SI Appendix*, Table S4). Finally, we found that some sOTUs, in particular from betacoronaviruses (e.g, u24667 and u175, both with humans among their hosts), have experienced frequent host switches, whereas others have not (e.g. u165, which is restricted to pigs). In particular, u944 (SARS-Cov-2) has experienced an intermediate number of host switches compared to other coronaviruses (*SI Appendix*, Fig. S9).

We also separately investigated the diversification dynamics of the two main clades of coronaviruses: the alpha- and betacoronaviruses. The undated version of ALE inferred, in alphacoronaviruses, 64 cospeciations, 0 duplication, 35 host switches, and 25 losses, as well as a recent origination in bats (69% of the reconciliations) and frequent intra-order host switches (80%). In contrast, in betacoronaviruses, a majority of the reconciliations (76%) had an origination in mammalian orders other than bats, including rodents (25%), Artiodactyla (18%), or Carnivora (13%), and involved inter-order host switches (61%). These results suggest that the ancestral coronavirus originated in bats, gave rise to alphacoronaviruses in bats, and switched to a different non-bat host where it evolved into betacoronaviruses. With 80 cospeciations, 0 duplication, 50 switches, and 36 losses, the fraction of host switches relative to cospeciation events is higher in betacoranaviruses (0.63) than in alphacoronaviruses (0.54). In both clades, host switches from bats still occurred preferentially toward humans or domesticated mammals, and >15% of the switches were time inconsistent. The dated version of ALE that forces host switches to be time-consistent failed to output a reconciliation for betacoronaviruses; in alphacoronaviruses, we obtained significant reconciliations with more “mismatch” events (67, including 38 host switches and 29 losses) than cospeciation events (65), suggesting cophylogenetic signal without phylogenetic congruence (Box 1; Fig 1; 36). While the latter result would be inconclusive, when it is combined with the numerous time-inconsistent switches found with the undated ALE version, it suggests that a scenario of codiversification is very unlikely.

### Sensitivity analyses

We carried a series of sensitivity analyses to assess the robustness of our analyses to potential biases or issues, summarized in Table S7.

We found qualitatively similar results when applying ALE on different sub-parts of the palmprint region, suggesting that the potential occurrence of recombination does not bias our conclusions (*SI Appendix*, Table S5). The percentage of originations inferred to occur in bats decreased in the analyses on the first sub-part, probably because using such a short fragment (75 aa-long) does not allow robust reconciliations. We also obtained consistent results using a reconciliation method based on maximum parsimony (eMPRess) instead of maximum likelihood (ALE). Whatever the costs that we set for the different reconciliation events, eMPRess estimated significant reconciliations (*P*<0.01). For instance, when favoring host switches, we inferred a recent origination in bats in 54% of the reconciliations and observed on average 32 cospeciations (s.d. +/-3), 2 losses (s.d. +/-1), 0.1 duplication (s.d. +/-0.3), and 140 host switches (s.d. +/-3) including several “back-in-time” host switches of >30 Myr. eMPRess therefore also supports a scenario of recent origination in bats and diversification by preferential host switches (Fig. 1B). Without investigating the time-consistency of the host switches, we would have wrongly concluded that almost one fourth of the diversification of Coronaviridae is related to ancient cospeciation events.

Our understanding of extant mammal-coronavirus interactions is incomplete and subject to taxonomic and geographic biases (43). To address potential biases, we investigated the potential effect of sampling biases on our results by sub-sampling our dataset. First, we tested the effect of unequal sampling effort within screened mammal species (e.g. humans and domesticated animals have been more extensively sampled than wild animals). When randomly subsampling only three Genbank accession codes per host species, we still found frequent originations in bats (59% +/- s.d. 5%), preferential host switches (48% +/- s.d. 5%), and frequent transfers from bats to humans (in 90% of the subsampled dataset) or domesticated animals (e.g. *Sus scrofa* and *Camelus dromedarius* in 100% and 52% of the subsampled dataset, respectively). Second, we tested the effect of unequal sampling effort across the mammal tree of life. When randomly subsampling up to 10 species per mammalian order, we observed an average decrease in the probability of origination in bats (from 56% in the original dataset to 37% +/- s.d. 10%), yet this scenario remained much more likely than an origination in other mammalian orders, such as artiodactyls (14% +/- s.d. 4%), rodents (11% +/- s.d. 3%), or carnivores (10% +/- s.d. 2%). Transfers from bats still occurred more frequently toward humans or domesticated mammals (*Camelus dromedarius*, *Rattus norvegicus*, or *Sus scrofa*): along with *Sorex araneus*, these species were in the top 5 of the species receiving a coronavirus from bats *ffing* . Overall, these sensitivity analyses indicate that the likely origination in bats and the frequent transfers from bats to humans or domesticated animals is not artifactually driven by the high number of bats species in the dataset nor the enhanced monitoring of coronaviruses in humans or domesticated animals.

### Geographical distribution of coronaviruses

An additional piece of evidence for a recent origination scenario comes from the geographical distribution of coronaviruses, with a hotspot of diversity in Eurasia that has not colonized the whole world (Fig. 5A, Fig. 1B). The coronavirus’ hotspot is more strongly influenced by the diversity of alphacoronaviruses than of betacoronaviruses (*SI Appendix*, Fig. S15). The higher host switches rates and broader host range of betacoronaviruses is reflected in a more widespread geographic distribution, with less pronounced hotspots when compared to alphacoronaviruses (*SI Appendix*, Fig. S15). Mammalian hosts of coronaviruses have a hotspot of species diversity concentrated in East Asia (Fig. 5C). The richness of coronaviruses presents a similar pattern, but with two comparable hotspots of species diversity in East Asia and Southern Europe (Fig. 5A), suggesting that the European hotspot is composed by fewer host species, together carrying as diverse a set of coronaviruses as the Asian hotspot. Other regions with a relatively high richness of coronaviruses and their hosts include parts of the African continent. The Americas and Australia have relatively low richness of coronaviruses and their hosts. Phylogenetic diversity of both hosts and coronaviruses (Fig. 5B, D) depict a similar pattern but with phylogenetic diversity more evenly distributed across most world regions, including the Americas.

**Figure 5.**
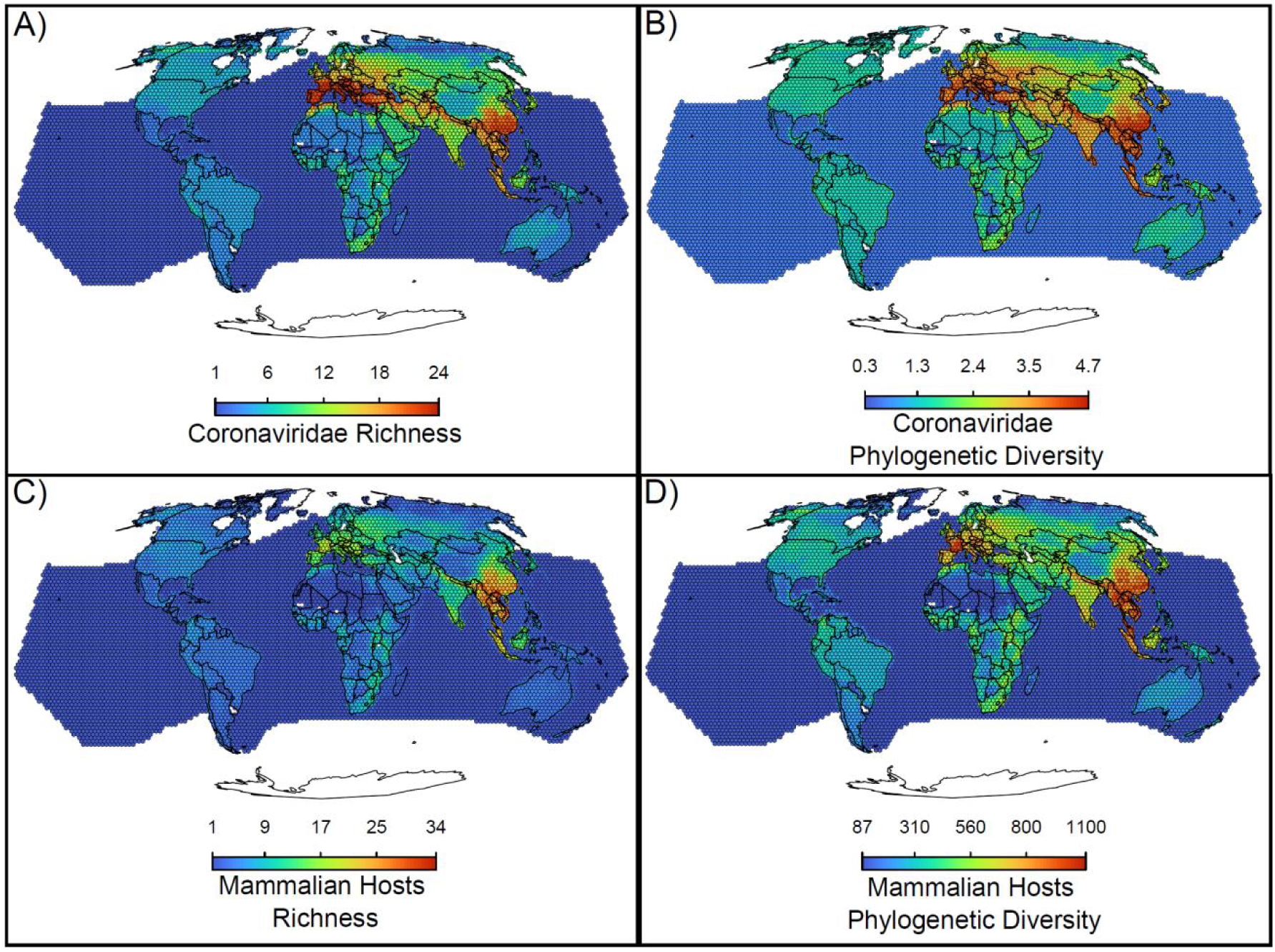
Maps of the diversity of coronaviruses and their mammal hosts. In A) the richness of species of coronaviruses; geographic range maps of coronaviruses were constructed after applying the host-filling method on the geographic range maps of mammalian hosts of coronaviruses. In B) Faith’s (1992) phylogenetic diversity of coronaviruses, calculated using the phylogenetic tree of coronaviruses (see main text). In C) and D), the richness and phylogenetic diversity of mammal hosts of coronaviruses, respectively. All maps are on the Mollweide projection.

## Discussion

### Inferring the Coronaviridae evolutionary history

Together, our results suggest that the common ancestor of extant mammalian coronaviruses originated recently in a bat species, and that coronaviruses diversification occurred via preferential host switches rather than through codiversification with mammals. Although we cannot unequivocally date the timing of origination of coronaviruses in mammals, we demonstrate that Coronaviridae is a highly dynamic clade in which diversification operates through host switches at a much faster pace than that of their hosts. sOTUs are rapidly replaced by newly-generated ones, with little role for codiversification with the hosts. The high diversity and endemicity of coronaviruses among bats have led others to anticipate that bats might be implicated in the origin of coronaviruses (2, 20, 23, 24), although definitive proof was lacking. We provided evidence for that hypothesis using a probabilistic cophylogenetic model and accounting for the entire known diversity of coronaviruses across mammals. Independent evidence for coronavirus recent host switches among different species exists in the literature (44, 45). The envisioned scenario suggests a timing of origination for extant Coronaviridae that is much more recent than the hundreds of millions of years ago suggested by (26). This is not surprising given the difficulties in estimating divergence times and inferring branch lengths for viral phylogenies (26, 27, 29), and provided that the dating of Wertheim et al. (26) relied on a substitution rate estimated from data with limited temporal signal (∼50 serially sampled contemporary sequences of a short gene fragment, (23)).

Our results contradict previous suggestions that codiversification with vertebrate hosts played an important role in Coronaviridae diversification (26, 28, 31). They also suggest that previously reported cases of long-term codiversification in vertebrate RNA viruses have been largely over-estimated, as many of them may instead be cases of diversification by host switches occurring preferentially among closely-related hosts (34). Indeed, these two scenarios both generate cophylogenetic signal in host-symbionts associations, such that cophylogenetic signal alone is not evidence for long-term codiversification (35). In addition, under a scenario of recent origination and preferential host switches, event-based cophylogenetic methods tend to artifactually favor biologically unrealistic scenarios with codiversification and back-in-time host switches, as we have shown here. As the time-consistency of host switches is typically not investigated, this has remained unnoticed, and evidence for codiversification has been taken for real. Ideally, cophylogenetic reconciliation methods would not allow such time-inconsistent host switches. However, imposing time constraints in methods based on parsimony is NP-hard (46), and the ‘dated’ version of ALE is not well adapted when recent host switch events dominate evolutionary history. We have found two ways to get around the problem, by interpreting time-inconsistent host switches as evidence for recent preferential host switches, and by gradually transforming the host tree to avoid large back-in-time switches, however future efforts should focus on developing time-consistent cophylogenetic methods. This would allow more robust and precise inferences of host-virus (and more generally host-symbiont) evolutionary history. We have shown by simulation that contrasting scenarios, such as codiversification, or diversification by preferential host switching after originating within or outside bats, leave distinct signatures in the data. This suggests that deep neural networks trained on simulated data, which have already shown their performance in phylodynamics (47, 48, 49, 50, could also be useful for the analysis of cophylogenetic data.

### Host switching dynamics in Coronaviridae

During their evolution, coronavirus’ host switches occurred more frequently within than between mammalian orders. This suggests that mammalian characteristics shared between relatives (e.g., genetic, behavioral, ecological), and the frequency of encounters among hosts play important roles in determining coronavirus’ host switches. Additionally, between-order host switches occurred more frequently among non-flying mammals and among orders containing humans and urban and domesticated mammals, suggesting that contact frequency alone is likely a key characteristic in host switches. Accordingly, amongst the most-likely host switches towards humans were those coming from mammals suspected to be involved in the transfer of specific coronavirus sOTUs likely through contact, for instance, camels in the case of MERS-CoV (44), Asian palm civets with SARS-CoV (16, 17) and the house mouse with SARS-CoV-2 (45). Importantly, we found that host switches from bats to other mammalian species were rare during the evolutionary history of Coronaviridae, even though coronaviruses originated and are more diverse within bats. These pieces of evidences suggest that bats are a closed reservoir of the Coronaviridae diversity, as also suggested by their relative isolation in the mammal-coronaviruses interaction network (Fig. 3).

Spillovers from bats to non-bat species, when they occurred, were found more likely to be towards humans than to any other mammalian species, suggesting humans may have acted as evolutionary intermediate hosts amongst mammals, in line with their centrality in the mammal-coronaviruses interaction network (Fig. 3). From an ecological perspective, the large abundance and widespread geographic distribution of humans, together with our habits of forcing contact with other species, including bats, make it unsurprising that humans, among all mammal species, have acted as intermediate hosts of ancestral forms of coronaviruses. Interestingly, for some individual species of coronaviruses, such as the SARS-CoV2 and other SARS-like coronaviruses, the dominant hypothesized scenario is that precursor forms spread from a bat to another intermediate mammalian host before infecting humans (20, 51). Our molecular marker lacks the intra-I resolution necessary to make species-level predictions, but our results suggest that more ancient coronaviruses host switches may have occurred in the other direction: from bats to humans to non-bat mammals. The large spillover of SARS-CoV2 from humans to wild mammalian lineages has now been well documented (51) and tend to confirm our results of humans as the intermediate host. Many human activities lend credit to the human-as-evolutionarily-intermediate-host-hypothesis, including human excursions to bat caves (52), hunting (53), and habitat destruction and modification (54), all of which increase the contact between bats and humans and their domesticated animals (54). Conservation of bats’ natural habitats, away from human contact, could help avoiding further spreads of coronaviruses among humans.

In agreement with previous studies (2, 21-24, we found noticeable differences between Alpha- and Betacoronaviruses. While Alphacoronaviruses likely originated in bats and mostly diversified by switching between bat lineages, betacoronaviruses most likely originated in a non-bat lineage and experienced a majority of their host switches between non-bat mammals. In addition, our findings suggest that host switching rates are higher in betacoronaviruses than in alphacoronaviruses. These different host switching dynamics are consistent with biogeographical differences: betacoronaviruses tend to be more geographically spread than alphacoronaviruses, which are more restricted to Eurasia. Some of the trends we observed (e.g. the higher host switching rates in betacoronaviruses) are not consistent with previous findings (55, 56, however these results are not directly comparable, as studies were conducted at very different scales (across the Americas (55), across China (56), and here at the global scale).

Insights of past and future host switches are gained from coronavirus geographic distribution. Coronaviruses are found worldwide and their hotspots of diversity are concentrated in East Asia and Southern Europe, where they likely originated. Previous assessments of the diversity of bat hosts of betacoronaviruses suggested similar hotspots but with a distribution of coronaviruses more concentrated in the hotspots (7, 14, 15) than the more pervasive pattern we found using all mammalian hosts. Moreover, the distribution of coronaviruses is less concentrated in the hotspots when phylogenetic metrics of diversity are included, suggesting that species richness alone is masking the global evolutionary potential of these viruses (57). Coronaviruses’ likely recent origination in bats, high within-order transmission rates, and their capacity to switch between mammal orders in some cases suggest the potential for future fast spreading and increase in the number of species across most world regions. Among alphacoronaviruses, the spread is more likely to remain concentrated within bats, while betacoronaviruses have a higher potential for among-orders spreading and infection of new mammalian hosts. The betacoronaviruses lineages already detected in humans are host generalists with high transmission rates, suggesting that continued monitoring may be wise in mitigating potential future pandemics.

### Limitations and perspectives

A few important limitations of our analyses deserve to be mentioned. First, we used a short marker gene to reconstruct the Coronaviridae phylogeny. However, the RdRp palmprint marker we used is very conserved and therefore routinely used for delimiting operational taxonomic units in RNA viruses and reconstructing their evolutionary history (38, 39). Our tree has an identical genus-level topology compared with trees constructed using other genome regions, such as the nucleocapsid portion of the coronavirus genome (23), and very close topologies were also observed using other genome regions such as the spike, envelope, and membrane regions (23), as well as the entire RdRp region (2, 6, 23): it differs only on the placement of Gammacoronavirus and Deltacoronavirus in relation to the others, but it is consistent in the monophyly of both Alphacoronavirus and Betacoronavirus and recovers the sister-clade relationship between Gamma and Delta. In addition, because the palmprint region is a conserved region, we could not reconstruct the recent evolutionary history of coronaviruses (i.e. the within sOTU transmission dynamic). Combining the palmprint region with a fast-evolving region(s) would enable more precise estimates of the recent routes of coronaviruses’ transmission, including that of SARS-CoV-2. More generally, characterizing multiple genetic regions with different evolutionary rates across Coronaviridae would allow us to more precisely elucidate the timescale of the evolutionary history of coronaviruses alongside their hosts.

Second, recombination is an important mechanism of viral evolution (58), and approaches more adequately designed to investigate the role of recombination are needed. The fact that different subparts of the palmprint region lead to similar results indicates that recombination acting on the palmprint region is unlikely to bias our conclusions. However, looking at other genomic regions would allow gaining a more complete understanding of the role of recombination in coronavirus evolution.

Third, the record of associations between coronaviruses and mammals is necessarily incomplete (not all mammal species have been screened for coronaviruses) and likely biased towards bats and mammals that are in contacts with humans. This a common bias when studying mammal-virus associations that may cause various issues (43.. However, our sub-sampling analyses suggest that our main results (recent origination in bats and frequent transfer from human-associated mammals to humans) are not artifactually driven by sampling biases. ALE explicitly accounts for undersampling by assuming that host switches involve unsampled intermediate hosts, which may explain the robustness of our findings to sampling biases.

### Concluding remarks

Understanding the evolutionary origins and diversification of viruses is useful for predicting new transmission routes, yet the relative frequencies of virus–host cospeciation versus cross-species transmission in the evolution of vertebrate RNA viruses remains uncertain (31). We found that coronaviruses originated in bats where they are more diverse nowadays, and later diversified in other mammal orders through preferential host switches. Spillovers from bats were rare but likely human-induced, suggesting humans are the intermediate evolutionary bridge that facilitated the spread of coronaviruses across mammals. Host switches between primates and artiodactyls, perissodactyls, and carnivorans occur frequently. This suggests a potential for the spread of coronaviruses to new mammalian hosts beyond their current prevalence in East Asia and Europe, and raises concerns about the possibility of future pandemics related to coronaviruses. Our results indicate that reducing human-bat contact, for instance through the conservation of bat habitats, could potentially serve as a mitigation strategy. They also suggest that cases of long-term virus–host codiversification, reported on the basis of cophylogenetic tests, have been largely over-estimated.

## Materials and Methods

### Operational Taxonomic Units for Coronaviridae

Viral species delimitation is difficult (59), and the number of proposed and/or estimated species or strains of Coronaviridae in the literature have varied (e.g., 100 proposed - 3,204 estimated (14); 88 proposed - 204 estimated (5)). However, an official assessment and classification of viruses are made by the International Committee on the Taxonomy of Viruses (ICTV - 60), by means of its Study Groups (59). The Coronaviridae Study Group of the ICTV (19) suggests that species delimitation within Coronaviridae should be made considering more than 90% amino acid sequence identity in conserved replicase domains as a criterion to include sequences in the same species (2). Estimates following ICTV suggestions have proposed between 17-39 species of coronaviruses (59; 61; 2), and the last (year 2021 v3) Master Species List from the ICTV lists 54 species in the family Coronaviridae (https://ictv.global/msl). Recently, a seminal paper by 38 proposed the use of a taxonomic barcode based on the palmprint region of the RdRp region for the systematic identification and classification of RNA viruses. The use of the RdRp region of the RNA genome is also common in other tree construction attempts for Coronaviridae (23; 26), given that the RdRp is an essential enzyme for replicating the RNA genome (62), and therefore aligns well with the ICTV proposition.

We therefore used the 46 described species-like Operational Taxonomic Units (sOTUs) for Coronaviridae delimited using ‘palmprint’ sequences by (38, 39). The palmprint is a conserved amino acid (aa) sub-sequence (150 aa in Coronaviridae) of central importance in the viral RdRp (38), selected for its homology across the large majority of sequences, allowing estimation of sequence divergence and phylogenetic trees (39). sOTUs were identified by Edgar et al. (38) after clustering palmprint sequences at 90% amino acid identity; and released through the Serratus project. Despite its relatively short length, trees constructed with this region are topologically equivalent to Coronaviridae trees based on other genes (see Discussion). IWe downloaded the palmprint amino acid sequences of Coronaviridae sOTUs from the Serratus project (https://serratus.io/; (38)) on April 13 of 2022.

### Mammalian hosts of Coronaviridae

All 46 sOTUs of Coronaviridae with a full palmprint and associated data in the NCBI database were screened for the identification of its hosts. From those, 35 sOTUs were associated with mammalian hosts and were kept for downstream analyses. Serratus’ associated metadata was used to identify GenBank accession codes linked to each sOTU. The complete set of 90,540 associated GenBank accession codes was screened to obtain the host information for each sOTU (on NCBI, Features>source>/host=). All the host species with a full Linnean name were kept as such. Accession codes with hosts leading to a generic level information were further inspected to identify the associated publication and determine the complete species name. Dubious cases or accession codes without publications had their hosts disregarded. Common names or high-level host information (e.g., host=“bats”) were generally eliminated except in a few cases where a domesticated species was found to be the host (i.e., host=“dog”,“canine” were *Canis lupus*; host=“cat”,“feline” were *Felis catus*; host=“pig”,“piglet”,“newborn piglet”,“sucking piglet”,“porcine”,“swine” were *Sus scrofa*). A final dataset of 116 mammalian hosts associated with the 35 sOTUs was assembled and used in downstream analyses. A matrix with the association between Coronaviridae sOTUs and mammalian species is available in *SI Appendix*, Dataset S1.

### Coronaviridae phylogenetic trees

We constructed a Coronaviridae tree using the palmprint amino acid sequence information of the 35 sOTUs. We aligned the amino acid sequences with MAFFT (63) and trimmed them with trimAl (64). The final alignment contained 150 amino acid positions. We used two main phylogenetic software, BEAST2 (65), which performs rooting and time calibration and PhyloBayes (66), which generates outputs adapted to the cophylogenetic algorithm we used. We visualized phylogenetic trees using R (67).

In order to run BEAST2, we generated an input file using BEAUti with 35 sOTU sequences and the following parameters: a WAG model with 4 classes of rates and invariant sites, a birth-death prior, and a relaxed log-normal clock. BEAST2 sampled a posterior distribution of ultrametric trees using Markov chain Monte Carlo (MCMC) with 4 independent chains each composed of 100,000,000 steps sampled every 10,000 generations. We checked the convergence of the 4 chains using Tracer (68). We used LogCombiner to merge the results setting a 25% burn-in and TreeAnnotator to obtain a Maximum Clade Credibility (MCC) tree with median branch lengths. Using a LG model instead of WAG also gave a consensus tree with a very similar topology.

To further assess the robustness of the BEAST2 tree rooting, we estimated the root position on a 46-sOTU maximum likelihood tree (from the Serratus pr–ject – (38)) assuming a strict molecular clock and an ultrametric tree. We used an ultrametric setting as temporal information from the tip dates (ranging between 1999 and 2022, a negligible difference with respect to the root age of dozens of thousands or even millions of years) was not sufficient to infer the mutation rate (we assessed the temporal signal with TempEst, (69)). We performed rooting and time-scaling with LSD2 (v2.3, (70)), assuming a tree of unknown scale (e.g. fixing all the tips dates to 1 and the root date to 0) with outlier removal and root search on all branches. LSD2 detected no outliers and positioned the root on the same branch as in the BEAST2 MCC tree (between alpha and betacoronaviruses).

The tree we constructed with PhyloBayes was specifically designed to the application of cophylogenetic methods, which cannot handle the presence of the same symbiont species in multiple host species. Following (37), we multiplicated the sOTU sequences in cases when a sOTU was present in several mammal species, such that each mammal-coronavirus association is represented by one single sequence (total of 173 sequences). We reconstructed the phylogenetic tree of the 173 sequences with PhyloBayes, run using an LG model, 4 classes of rates, and a chain composed of 4,000 steps with a 25% burn-in. To sum up, we built two trees for coronaviruses: a 35-tip tree with BEAST2 (with one tip per sOTU) and a 173-tip tree with PhyloBayes (with multiple tips per sOTU corresponding to the multiple host species associated with each sOTU, such that cophylogenetic methods can run).

### Mammalian phylogenetic tree

We obtained a phylogenetic hypothesis for mammals from the consensus DNA-only tree of (71), one of the most complete and updated phylogenies for mammals. We downloaded the node-dated tree for 4,098 mammals, constructed based on a 31-gene supermatrix, from the VertLife website (http://vertlife.org/data/mammals/). We used a pruned version of the tree with the 116 mammalian hosts of Coronaviridae in all analyses in this paper. We kept for each node the 95% credible interval of its age estimate.

### Phylogenetic signal in the association between coronaviruses and mammals

To assess whether closely related coronaviruses interact with similar mammals, and vice-versa, i.e. presence of phylogenetic signal in the association, we used Mantel tests following (40). Mantel tests were constructed by taking the Pearson correlation between phylogenetic distances and ecological distances. Phylogenetic distances of coronaviruses were computed on the BEAST2 MCC phylogeny. Ecological distances were calculated based on the interaction network matrix containing the association between coronavirus’ sOTUs and mammals, accounting for the evolutionary relationships among interaction partners using UniFrac distances (72). Firstly, we conducted Mantel tests permuting the identity of species but keeping the number of partners per species constant; this allows for assessing the effect of species identity while controlling for the confounding effect of the number of partners. Then, we evaluated the phylogenetic signal in the number of partners alone. Lastly, we calculated clade-specific Mantel tests for sub-networks containing at least 10 species (40) to evaluate whether phylogenetic signal was stronger for specific subclades of mammals or coronaviruses. Ten thousand permutations were used in each analysis to assess significance. Analyses were conducted using the phylosignal_network and phylosignal_sub_network functions in the R package RPANDA (73).

### Coronaviridae origination and host switches

We used the amalgamated likelihood estimation–(ALE – (41)) to reconcile the mammal and coronaviruses evolutionary history using events of cospeciations, host switches, duplications, and losses. Originally designed in the context of gene tree – species tree reconciliations (41), ALE has also been particularly useful in the context of host-symbiont cophylogenetic analyses as it considers both phylogenetic uncertainty of the symbiont evolutionary history and undersampling of host species (37, 74, 75). ALE indeed assumes that host switches may imply an unsampled or extinct intermediate host lineage (42). ALE therefore intrinsically accounts for the incompleteness of our dataset, *i.e.* the fact that we only observe a subsample of mammalian species for which coronaviruses have been detected among all the infected species.

We ran ALE with the posterior distribution of phylogenetic trees of coronaviruses generated with PhyloBayes to estimate the maximum likelihood rates of host switches, duplications, and losses of the coronaviruses. We first tried running the “dated” version of ALE, which accounts for the order of branching events in the host phylogeny, therefore only allowing for time-consistent host switches (i.e. host switches that happen between two contemporary host lineages). However, this led to unrealistic parameter estimates (such as very high loss rates) and ALE was not able to output possible reconciliations, suggesting that the mammalian and Coronaviridae trees are too incongruent to be reconciled with only time-consistent host switches, *i.e.* that the scenario of codiversification is likely unrealistic. We therefore used the “undated” version of ALE that only exploits the topology of both the host and the symbiont tree and thus does not constrain the host switches to be time-consistent. ALE generated a total of 5,000 reconciliations, from which we extracted the mean number of cospeciations, host switches, duplications, and losses. We also reported the likely origination of coronaviruses in mammals (i.e. the branch in the mammal phylogeny that was first infected by coronaviruses) by computing, for each branch of the mammalian tree, the frequency of reconciliations (among the 5,000) that supported an origination in that branch. If a reconciliation requires more cospeciation events and fewer host switch events, than expected under a null scenario of independent evolution, this indicates that the evolution of the symbiont was not independent of that of the host, and in this case, we talk about a “significant reconciliation” (76).

We evaluated the significance of the reconciliation by comparing the estimated number of cospeciation and host switch events to null expectations obtained with ALE by shuffling the mammal host species across the mammal tree, both randomly or within major biogeographic regions according to the proposal of regions by (77) for mammals (six biogeographic regions: North American, South American, African, Eurasian, Oriental, and Australian). We considered a reconciliation to be significant if the observed number of cospeciations was higher than 95% of the null expectations and if the number of host switches was lower than 95% of the null expectations (37). The likeliness of a host switch between two mammal lineages is measured as the frequency of the reconciliations in which it occurs. Finally, we reported the ratio of time-inconsistent host switches by focusing on “back-in-time” switches, from a donor mammal lineage to an older receiver mammal lineage that never coexisted. We identified time-inconsistent host switches directly on the consensus mammalian phylogeny, or considering the 95% credible interval around each age node estimate to avoid counting time-inconsistent host switches that may arise from incorrect estimates of divergence times.

Because ALE estimated a large proportion of time-inconsistent host switches (see Results), we first tested the scenario of a more recent origination by collapsing all mammalian nodes anterior to X Myr into a polytomy at the root of the phylogeny (with X varying from 55 Myr to 5 Myr), such that the coronavirus origination and host switches inferred by ALE could not involve mammal lineages older than X Myr. Second, we investigated the scenario of diversification by pure preferential host switches of the coronaviruses among extant mammals. To do so, we ran ALE on a star mammalian phylogenetic tree. In this context, ALE could no longer infer cospeciations, and only fit events of host switches, duplications, or losses. When inferring a likely host switch between two specific mammalian lineages on a star phylogeny, there are often as many reconciliations suggesting one directionality of the host switch (i.e. from one of the lineages to the other) as the other. We then only kept host switches present in at least 10% of the reconciliations and looked at the ratio between the number of host switches that were estimated within versus between mammal orders. We compared this ratio to a null expectation obtained by randomly shuffling the host mammal species.

We conducted the same set of analyses on the sub-datasets formed by alpha- and betacoronaviruses separately. The dated version of ALE was able to output reconciliations for the alphacoronaviruses dataset, but not for the betacoronaviruses.

Recombination is frequent in viruses and the palmprint region may be recombined, such that different fragments of the palmprint region may have different evolutionary histories, potentially biasing our inference. We carried several recombination tests (OpenRDP https://github.com/PoonLab/OpenRDP, our own custom code, and Gubbins, 78 that were inconclusive, suggesting that the palmprint region is too short to infer anything about recombination. We therefore instead tested whether the results we obtained on the whole 150-amino acid palmprint region could be impacted by recombination by replicating the ALE analyses on two sub-regions: the first part (positions 1-75) and the last part (positions 76-150).

We also repeated our cophylogenetic analyses using eMPRess (46), another event-based cophylogenetic approach that reconciles host-symbiont evolutionary histories using maximum parsimony. eMPRess is a recent improved version of the popular Jane approach (32); it differs from Jane especially by not relying only on a heuristic and therefore guarantying that the solution truly corresponds to the maximum parsimony reconciliation(s) (46). However, contrary to Jane, eMPRess does not offer the possibility to constrain host switches to occur only among lineages from pre-specified time periods. eMPRess requires specifying cost values for the events of host switches (t), duplications (d), and losses (l). We tested two sets of cost values: (1) cost values that disadvantage host switches (d=6, t=6, l=1) and (2) uniform cost values that favor host switches (d=1, t=1, l=1). As with ALE, we evaluated the significance of the reconciliations using permutations. We ran eMPRess analyses on a set of 50 trees randomly sampled from the posterior distribution of PhyloBayes.

### Sampling biases

More effort has been put on the characterization of coronaviruses associated with humans, domesticated animals, and bats (bats represent 47% of the mammalian species in our dataset, in part because many bat species have been intensively screened for viruses). This can lead to two main sampling biases that could potentially impact our conclusions: (i) unequal sampling effort across mammalian species that have been screened, and (ii) unequal screening of mammalian species across the mammal tree of life. To assert whether such sampling biases could generate spurious results we designed two subsampling strategies and re-run the PhyloBayes and ALE analyses on each subsampled dataset. First, For the unequal sampling effort across mammalian species, we randomly subsampled only 3 Genbank accession codes per host species (and removed species described by less than 3 accessions). We replicated the subsampling 50 times and ran ALE on each subsampled dataset. The resulting dataset contained a total of 46 host species. Second, for the unequal screening of mammalian species across the mammal tree of life, we subsampled the dataset at the level of mammalian orders. Thus, we randomly subsampled up to 10 species per mammalian order. Only 4 mammalian orders (Chiroptera, Rodentia, Artiodactyla, and Carnivora) had at least 10 species, the other orders were kept unmodified. We replicated the subsampling 50 times and ran ALE on each subsampled dataset. The resulting dataset contained a total of 53 mammalian species.

### Simulation analyses

By running the undated version of ALE either on the mammal phylogeny or a star phylogeny, we proposed a framework to evaluate whether the cophylogenetic pattern is due to a history of ancient codiversification (i.e. a mix of cospeciations, host switches, duplications, and losses; Fig. 1A) or to a scenario where the coronaviruses diversify more recently by preferential host switches ((35); Fig. 1B). To validate the interpretation of our ALE results, we performed simulations under the two alternative scenarios of codiversification and diversification by preferential host switches. For the scenario of codiversification, we assumed that coronaviruses originated in the ancestors of bats and that they subsequently codiversified with the mammals by experiencing events of cospeciations, host switches, duplications, and losses. We used the function *sim_microbiota* in the R-package HOME to obtain the corresponding coronavirus sequences and coronavirus-mammals associations (79). We sampled a number of host switches uniformly between 100 and 150, used a duplication rate of 0.001, and simulated losses by host switching replacement. Given the simulated coronaviruses phylogenies, we simulated on it the evolution of a 450-bp nucleotide sequence using a K80 model with a substitution rate of 0.5 and an expected proportion of variable sites of 50% (see 37, for further details on the model).

For the scenario of coronaviruses diversification by preferential host switches, we used a birth-death model (pbtree function in the R-package phytools) to simulate a phylogenetic tree of the coronaviruses: in our model, each coronavirus lineage is associated with a single host species, a birth event corresponds to a host switch (at rate 50), while a death event corresponds to a loss of a coronavirus in a host lineage (at rate 5). We started the diversification by assuming a single coronavirus infection in *Eidolon helvum* (a bat host of external lineages within betacoroviruses, u25738 and u27845). Then, following de (80) and (81), we modeled preferential host switches by assuming that for a host switch from a given donor mammal species, each potential receiver species has a probability proportional to exp(-0.035\**d*) where *d* is the phylogenetic distance between the donor and receiver species. Finally, we simulated RNA sequences of the coronavirus sequences using the function *simulate_alignment* in HOME. For each type of simulation, we generated 50 simulated datasets of mammal-coronavirus associations. For each dataset, we ran PhyloBayes and ALE on both the mammalian phylogeny and the star phylogeny.

In addition, we used simulations to test for a scenario where coronaviruses originated outside of bats but diversify faster within-bats than within other mammalian lineages, as previously suggested given the efficient immune systems of bats (82). We simulated originations in rodents followed by diversification by preferential host switches (as above) with a host switch rate twice more important in bats (rate 80) than in other mammals (rate 40). For each simulated dataset, we ran PhyloBayes and ALE on the star phylogeny and reported the percentage of originations incorrectly inferred in bats. Parameters of the simulations were chosen to mimick the diversity of the original mammal-coronavirus associations.

### Geographic distribution of Coronaviridae

We downloaded geographic range maps for each mammalian host species, with the exception of *Homo sapiens*, from the Map of Life website (https://mol.org/species/); see (83). These maps follow the taxonomy of the Mammal Diversity Database (84) supplemented with the Handbook of the Mammals of the World (HMW) database and the Alien Checklist database for invasive species (83).

We created a world map with hexagonal, equal-area grid cells of 220km on which we mapped host and coronavirus species diversity, using the Mollweide world projection to accurately represent areas. At large spatial scales, cells with ∼220km resolution return more reliable diversity estimates than smaller cells (85). We considered that a host species was present in any given cell if its range covered at least 30% of the cell area to avoid overestimating diversity. We calculated host species diversity as a simple sum of the species occurring in any given cell, and host phylogenetic diversity as Faith’s phylogenetic diversity inde– (PD – (86)) for each cell. We mapped Coronaviridae diversity using the host-filling method (87): we constructed a range map for each Coronaviridae sOTU by overlapping the range maps of all its hosts. We consider the host filling method appropriate in this case because coronaviruses are obligatory parasites that can only live inside hosts. Next, we calculated Coronaviridae sOTU diversity by summing range maps overlapping on each cell, and Coronaviridae phylogenetic diversity as Faith’s PD (86). We created these maps in R (67) using the packages epm (88), sf (89), and ape (90).

## Acknowledgments

This work was performed using HPC resources from GENCI-IDRIS (Grants 2021-A0100312405 and 2022-AD010313735). The authors thank B. Boussau and the Morlon lab for helpful discussions. RM thanks Campus France and the One Health/Make Our Planet Again Program for a fellowship to conduct this research.

## References

1. K. P. Alekseev, et al., Bovine-Like Coronaviruses Isolated from Four Species of Captive Wild Ruminants Are Homologous to Bovine Coronaviruses, Based on Complete Genomic Sequences. J Virol 82, 12422–12431 (2008).

2. R. J. De Groot, et al., Family : Coronaviridae. ICTV Ninth Report. https://ictv.global/report_9th/RNApos/Nidovirales/Coronaviridae (2022).

3. A. M. Q. King, Adams M.J., Carstens E.B., Lefkowitz E.J., Virus taxonomy: classification and nomenclature of viruses: Ninth Report of the International Committee on Taxonomy of Viruses (Elsevier, 2012).

4. J. C. Leao, et al., Coronaviridae—Old friends, new enemy! Oral Dis 28, 858–866 (2022).

5. M. Wardeh, M. Baylis, M. S. C. Blagrove, Predicting mammalian hosts in which novel coronaviruses can be generated. Nat Commun 12 (2021).

6. C. M. Zmasek, E. J. Lefkowitz, A. Niewiadomska, R. H. Scheuermann, Genomic evolution of the Coronaviridae family. Virology 570, 123–133 (2022).

7. D. J. Becker, et al., Optimising predictive models to prioritise viral discovery in zoonotic reservoirs. Lancet Microbe 3, e625–e637 (2022).

8. C. Drosten, et al., Identification of a Novel Coronavirus in Patients with Severe Acute Respiratory Syndrome. New England Journal of Medicine 348, 1967–1976 (2003).

9. J. Peiris, et al., Coronavirus as a possible cause of severe acute respiratory syndrome. The Lancet 361, 1319–1325 (2003).

10. A. M. Zaki, S. van Boheemen, T. M. Bestebroer, A. D. M. E. Osterhaus, R. A. M. Fouchier, Isolation of a Novel Coronavirus from a Man with Pneumonia in Saudi Arabia. New England Journal of Medicine 367, 1814–1820 (2012).

11. P. Zhou, et al., A pneumonia outbreak associated with a new coronavirus of probable bat origin. Nature 579, 270–273 (2020).

12. H. Ritchie, et al., Coronavirus Pandemic (COVID-19). Published online at OurWorldInData.org (2024).

13. P. C. Y. Woo, Y. Huang, S. K. P. Lau, K. Y. Yuen, Coronavirus genomics and bioinformatics analysis. Viruses 2, 1805–1820 (2010).

14. S. J. Anthony, et al., Global patterns in coronavirus diversity. Virus Evol 3 (2017).

15. N. F. R. Munoz, et al., The coevolutionary mosaic of bat betacoronavirus emergence risk. EcoEvoRXiv (2022).

16. Y. Guan, et al., Isolation and characterization of viruses related to the SARS coronavirus from animals in Southern China. Science (1979) 302, 276–278 (2003).

17. C. SMEC, Molecular Evolution of the SARS Coronavirus During the Course of the SARS Epidemic in China. Science (1979) 303, 1666–1669 (2004).

18. R. L. Graham, R. S. Baric, Recombination, Reservoirs, and the Modular Spike: Mechanisms of Coronavirus Cross-Species Transmission. J Virol 84, 3134–3146 (2010).

19. R. J. de Groot, et al., Commentary: Middle East Respiratory Syndrome Coronavirus (MERS-CoV): Announcement of the Coronavirus Study Group. J Virol 87, 7790–7792 (2013).

20. V. M. Corman, D. Muth, D. Niemeyer, C. Drosten, “Hosts and Sources of Endemic Human Coronaviruses” in Advances in Virus Research, (Academic Press Inc., 2018), pp. 163–188.

21. E. Mavrodiev, M. L. Tursky, S. Vincent, D. M. Williams, L. Schroder, On Classication and Taxonomy of Coronaviruses (Riboviria, Nidovirales, Coronaviridae) with Special Focus on Severe Acute Respiratory Syndrome-Related Coronavirus 2 (SARS-CoV-2) (2021) 10.21203/rs.3.rs-282371/v1.

22. P. C. Y. Woo, S. K. P. Lau, Y. Huang, K. Y. Yuen, Coronavirus diversity, phylogeny and interspecies jumping. Exp Biol Med 234, 1117–1127 (2009).

23. P. C. Y. Woo, et al., Discovery of Seven Novel Mammalian and Avian Coronaviruses in the Genus Deltacoronavirus Supports Bat Coronaviruses as the Gene Source of Alphacoronavirus and Betacoronavirus and Avian Coronaviruses as the Gene Source of Gammacoronavirus and Deltacoronavirus. J Virol 86, 3995–4008 (2012).

24. D. Vijaykrishna, et al., Evolutionary Insights into the Ecology of Coronaviruses. J Virol 81, 4012–4020 (2007).

25. A. J. Drummond, S. Y. W. Ho, M. J. Phillips, A. Rambaut, Relaxed Phylogenetics and Dating with Confidence. PloS Biol 4, e88 (2006).

26. J. O. Wertheim, D. K. W. Chu, J. S. M. Peiris, S. L. Kosakovsky Pond, L. L. M. Poon, A Case for the Ancient Origin of Coronaviruses. J Virol 87, 7039–7045 (2013).

27. J. O. Wertheim, S. L. Kosakovsky Pond, Purifying selection can obscure the ancient age of viral lineages. Mol Biol Evol 28, 3355–3365 (2011).

28. D. T. S. Hayman, M. A. Knox, Estimating the age of the subfamily Orthocoronavirinae using host divergence times as calibration ages at two internal nodes. Virology 563, 20–27 (2021).

29. S. Duchêne, E. C. Holmes, S. Y. W. Ho, Analyses of evolutionary dynamics in viruses are hindered by a time-dependent bias in rate estimates. Proceedings of the Royal Society B: Biological Sciences 281, 20140732 (2014).

30. A. Katzourakis, R. J. Gifford, Endogenous Viral Elements in Animal Genomes. PloS Genet 6, e1001191 (2010).

31. M. Shi, et al., The evolutionary history of vertebrate RNA viruses. Nature 556, 197–202 (2018).

32. C. Conow, D. Fielder, Y. Ovadia, R. Libeskind-Hadas, Jane: A new tool for the cophylogeny reconstruction problem. Algorithms for Molecular Biology 5 (2010).

33. Y.-Z. Zhang, W.-C. Wu, M. Shi, E. C. Holmes, The diversity, evolution and origins of vertebrate RNA viruses. Curr Opin Virol 31, 9–16 (2018).

34. J. L. Geoghegan, S. Duchêne, E. C. Holmes, Comparative analysis estimates the relative frequencies of co-divergence and cross-species transmission within viral families. Plos Pathogens 13(2): e1006215 (2017).

35. D. M. de Vienne, et al., Cospeciation vs host-shift speciation: methods for testing, evidence from natural associations and relation to coevolution. New Phytologist 198, 347– 385 (2013).

36. B. Perez-Lamarque, H. Morlon, Distinguishing cophylogenetic signal from phylogenetic congruence clarifies the interplay between evolutionary history and species interactions. Syst Biol, 13 (2024) 10.1093/sysbio/syae013.

37. B. Perez-Lamarque, H. Morlon, Comparing different computational approaches for detecting long-term vertical transmission in host-associated microbiota in Molecular Ecology, (John Wiley and Sons Inc, 2022) 10.1111/mec.16681.

38. R. C. Edgar, et al., Petabase-scale sequence alignment catalyses viral discovery. Nature 602, 142–147 (2022).

39. A. Babaian, R. Edgar, Ribovirus classification by a polymerase barcode sequence. PeerJ 10 (2022).

40. B. Perez-Lamarque, et al., Do closely related species interact with similar partners? Testing for phylogenetic signal in bipartite interaction networks. Peer Community Journal 2, article no. e59 (2022).

41. G. J. Szöllosi, W. Rosikiewicz, B. Boussau, E. Tannier, V. Daubin, Efficient exploration of the space of reconciled gene trees. Syst Biol 62, 901–912 (2013).

42. G. J. Szöllosi, E. Tannier, N. Lartillot, V. Daubin, Lateral gene transfer from the dead. Syst Biol 62, 386–397 (2013).

43. T. Poisot, et al., Network embedding unveils the hidden interactions in the mammalian virome. Patterns 4, 100738 (2023).

44. G. Dudas, L. M. Carvalho, A. Rambaut, T. Bedford, MERS-CoV spillover at the camel-human interface. Elife 7 (2018).

45. C. Wei, et al., Evidence for a mouse origin of the SARS-CoV-2 Omicron variant. Journal of Genetics and Genomics 48, 1111–1121 (2021).

46. S. Santichaivekin, et al., eMPRess: a systematic cophylogeny reconciliation tool. Bioinformatics 37, 2481–2482 (2021).

47. I. Lajaaiti, S. Lambert, J. Voznica, H. Morlon, F. Hartig, A comparison of deep learning architectures for inferring parameters of diversification models from extant phylogenies. BioRxiv 2023.03.03.530992 (2023). 10.1101/2023.03.03.530992.

48. A. Thompson, B. Liebeskind, E. J. Scully, M. Landis, Deep learning and likelihood approaches for viral phylogeography converge on the same answers whether the inference model is right or wrong. BioRxiv 2023.02.08.527714 (2023). 10.1101/2023.02.08.527714.

49. J. Voznica, A. Zhukova, V. Boskova, E. Saulnier, F. Lemoine, M. Moslonka-Lefebvre, O. Gascuel, Deep learning from phylogenies to uncover the epidemiological dynamics of outbreaks. Nat Commun 13, 3896 (2022).

50. S. Lambert, J. Voznica, H. Morlon, Deep learning from phylogenies for diversification analyses. Syst Biol, 72, 1262–1279 (2023).

51 C. C. S. Tan, et al., Transmission of SARS-CoV-2 from humans to animals and potential host adaptation. Nat Commun 13 (2022).

52. 52. N. M. Furey, P. A. Racey, “Conservation Ecology of Cave Bats” in Bats in the Anthropocene: Conservation of Bats in a Changing World, (Springer International Publishing, 2016), pp. 463–500.

53. 53. T. Mildenstein, I. Tanshi, P. A. Racey, “Exploitation of Bats for Bushmeat and Medicine” in Bats in the Anthropocene: Conservation of Bats in a Changing World, (Springer International Publishing, 2016), pp. 325–375.

54. I. Smith, L. F. Wang, Bats and their virome: An important source of emerging viruses capable of infecting humans. Curr Opin Virol 3, 84–91 (2013).

55. D. A. Caraballo, Cross-species transmission of bat coronaviruses in the Americas: contrasting patterns between alphacoronavirus and betacoronavirus. Microbiology Spectrum, 10, 4, e01411–22 (2022).

56. A. Latinne, et al., Origin and cross-species transmission of bat coronaviruses in China. Nat Comm 11, 4235 (2020).

57. S. Leopardi, et al., Interplay between co-divergence and cross-species transmission in the evolutionary history of bat coronaviruses. Infection, Genetics and Evolution 58, 279– 289 (2018).

58. M. Pérez-Losada, M. Arenas, J. C. Galán, F. Palero, F. González-Candelas, Recombination in viruses: Mechanisms, methods of study, and evolutionary consequences. Infection, Genetics and Evolution 30, 296–307 (2015).

59. A. E. Gorbalenya, et al. The species Severe acute respiratory syndrome-related coronavirus: classifying 2019-nCoV and naming it SARS-CoV-2. Nature Microbiology, 5(4), 536–544 (2020). 10.1038/s41564-020-0695-z.

60. M. J. Adams, et al. 50 years of the International Committee on Taxonomy of Viruses: progress and prospects. Arch. Virol. 162, 1441–1446 (2017).

61. F. S. Campos, R Lourenço-de-Moraes, Ecological Fever: The Evolutionary History of Coronavirus in Human-Wildlife Relationships. Front. Ecol. Evol. 8:575286. (2020) doi: 10.3389/fevo.2020.575286

62. C. Venkataraman, M. Brauer, K. Tibrewal, P. Sadavarte, Q. Ma, A. Cohen, S. Chaliyakunnel, J. Frostad, Z. Klimont, R.V. Martin, D.B. Millet, S. Philip, K. Walker, S. Wang, Source influence on emission pathways and ambient PM2.5 pollution over India (2015–2050) Atmos. Chem. Phys., 18, 8017–8039 (2018).

63. K. Katoh, D. M. Standley, MAFFT Multiple Sequence Alignment Software Version 7: Improvements in Performance and Usability. Mol Biol Evol 30, 772–780 (2013).

64. S. Capella-Gutiérrez, J. M. Silla-Martínez, T. Gabaldón, trimAl: a tool for automated alignment trimming in large-scale phylogenetic analyses. Bioinformatics 25, 1972–1973 (2009).

65. R. Bouckaert, et al., BEAST 2: A Software Platform for Bayesian Evolutionary Analysis. PLoS Comput Biol 10, e1003537 (2014).

66. N. Lartillot, H. Philippe, A Bayesian Mixture Model for Across-Site Heterogeneities in the Amino-Acid Replacement Process. Mol Biol Evol 21, 1095–1109 (2004).

67. R Core Team, R: a language and environment for statistical computing (2018).

68. A. Rambaut, A. J. Drummond, D. Xie, G. Baele, M. A. Suchard, Posterior Summarization in Bayesian Phylogenetics Using Tracer 1.7. Syst Biol 67, 901–904 (2018).

69. A. Rambaut, T. T. Lam, L. Max Carvalho, O. G. Pybus, Exploring the temporal structure of heterochronous sequences using TempEst (formerly Path-O-Gen). Virus Evol 2, vew007 (2016).

70. T.-H. To, M. Jung, S. Lycett, O. Gascuel, Fast Dating Using Least-Squares Criteria and Algorithms. Syst Biol 65, 82–97 (2016).

71. N. S. Upham, J. A. Esselstyn, W. Jetz, Inferring the mammal tree: Species-level sets of phylogenies for questions in ecology, evolution, and conservation. PLoS Biol 17 (2019).

72. J. Chen, et al., Associating microbiome composition with environmental covariates using generalized UniFrac distances. Bioinformatics 28, 2106–2113 (2012).

73. H. Morlon, et al., RPANDA: an R package for macroevolutionary analyses on phylogenetic trees. Methods Ecol Evol 7, 589–597 (2016).

74. M. Groussin, et al., Unraveling the processes shaping mammalian gut microbiomes over evolutionary time. Nat Commun 8 (2017).

75. M. Bailly-Bechet, et al., How Long Does Wolbachia Remain on Board? Mol Biol Evol 34, 1183–1193 (2017).

76. R. G. Dorrell, et al., Phylogenomic fingerprinting of tempo and functions of horizontal gene transfer within ochrophytes 10.1073/pnas.2009974118/-/DCSupplemental.

77. C. Barry Cox, The biogeographic regions reconsidered. J Biogeogr 28, 511–523 (2001).

78. N. J. Croucher, et al, Rapid phylogenetic analysis of large samples of recombinant bacterial whole genome sequences using Gubbins, Nucleic Acids Research, 43, 3, e15 (2014).

79. B. Perez-Lamarque, H. Morlon, Characterizing symbiont inheritance during host– microbiota evolution: Application to the great apes gut microbiota. Mol Ecol Resour 19, 1659–1671 (2019).

80. D. M., de Vienne, T. Giraud, J. A. Shykoff, When can host shifts produce congruent host and parasite phylogenies? A simulation approach. J Evol Biol 20, 1428–1438 (2007).

81. B. Perez-Lamarque, H. Krehenwinkel, R. G. Gillespie, H. Morlon, Limited evidence for microbial transmission in the phylosymbiosis between Hawaiian spiders and their microbiota. mSystems 7, e01104–21 (2022).

82. A. Banerjee, et al., Novel insights into immune systems of bats. Front. Immunol. 11, 10.3389/fimmu.2020.00026 (2020).

83. C. J. Marsh, et al., Expert range maps of global mammal distributions harmonised to three taxonomic authorities. J Biogeogr 49, 979–992 (2022).

84. C. J. Burgin, J. P. Colella, P. L. Kahn, N. S. Upham, How many species of mammals are there? J Mammal 99, 1–14 (2018).

85. A. H. Hurlbert, W. Jetz, Species richness, hotspots, and the scale dependence of range maps in ecology and conservation. PNAS 104, 13384–13389 (2007).

86. D. P. Faith, Conservation evaluation and phylogenetic diversity. Biol Cons 61, 1–10 (1992).

87. P. Pappalardo, et al., Comparing methods for mapping global parasite diversity. Global Ecol Biogeogr 29, 182–193 (2019).

88. P. O. Title, D. L. Swiderski, M. L. Zelditch, EcoPhyloMapper: an R package for integrating geographical ranges, phylogeny and morphology. Methods Ecol Evol 13, 1912–1922 (2022).

89. E. Pebesma, Simple features for R: standardized support for spatial vector data. The R Journal 10, 439–446 (2018).

90. E. Paradis, K. Schliep, ape 5.0: an environment for modern phylogenetics and evolutionary analyses in R. Bioinformatics 35, 526–528 (2019).

