## Supplementary material for "Recent evolutionary origin and localized diversity hotspots of mammalian coronaviruses": SI Appendix

**This PDF file includes:**

Box S1
Figures S1 to S15
Tables S1 to S7

**Other supporting materials for this manuscript include the following:**

Datasets S1

**Box S1:** Definitions.

Our terminology is as in Perez-Lamarque & Morlon (2024, Systematic Biology, doi: 10.1093/sysbio/syae013).

**Codiversification/Co-cladogenesis/Co-divergence:** Pattern of concomitant diversification events happening in both host and symbiont clades. Codiversification can occur due to processes of phylogenetic tracking or successive vicariance events affecting both clades.

**Coevolution:** Process of reciprocal evolutionary changes induced by selective pressures in two (or more) interacting lineages.

**Cophylogenetic signal:** Pattern depicting the tendency of closely related species to interact with closely related partners.

**Cospeciation:** Concomitant event of host and symbiont speciations.

**Event-based methods:** Cophylogenetic methods reconciling the host and symbiont phylogenies by fitting reconciliation events (e.g. cospeciation, host transfer, duplication, or loss) on the symbiont phylogeny.

**Phylogenetic congruence:** Pattern of high similarity of the phylogenetic trees of interacting host and symbiont clades in terms of topology and relative branch lengths. If host and symbiont divergence times are matching, phylogenetic congruence can correspond to codiversification.

**Phylogenetic signal:** Pattern depicting the tendency of closely related species to have similar traits.

**Diversification by preferential host switching:** Tendency of symbionts to experience host transfers toward closely related host species; when the transfer results in a speciation event in the symbiont lineage, this tends to generate phylogenetic congruence. This does not imply co-diversification though, as the divergence times of the symbionts may be much more recent than those of the hosts.

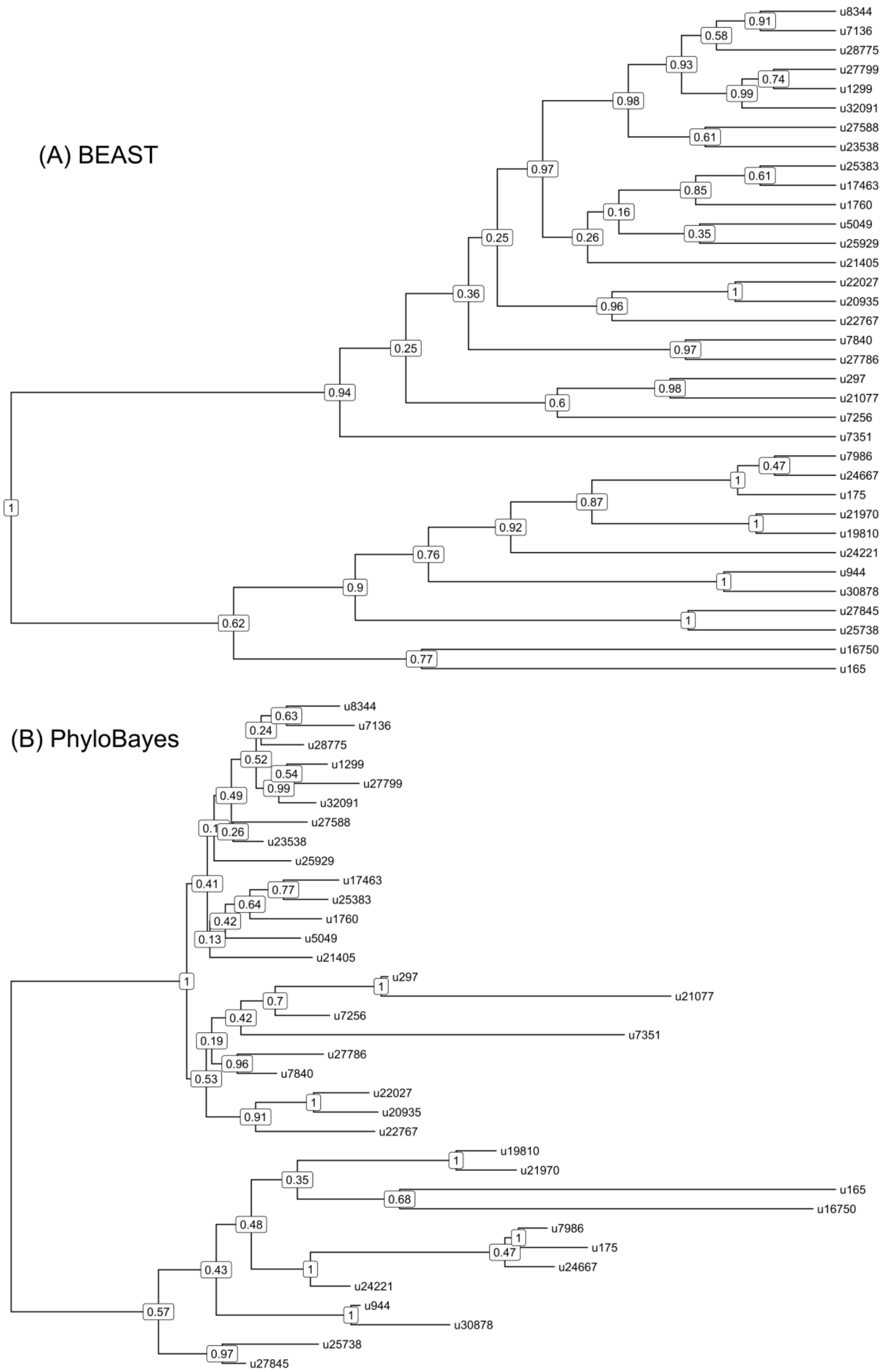


**Fig. S1. Phylogenetic relationships among coronaviruses sOTUs.** Consensus Coronaviridae tree constructed in (A) BEAST2 and (B) PhyloBayes using the palmprint amino acid sequence information of the 35 sOTUs of coronaviruses infecting mammals. We pruned out multiple sequences per sOTU in the PhyloBayes tree (that we included for reconciliation analyses) to represent a OTU-level tree comparable the one obtained with BEAST2. u16750 corresponds to gammacoronaviruses, and u165 to deltacoronaviruses; the top subtrees correspond to alphacoronaviruses while the bottom subtrees correspond to betacoronaviruses.

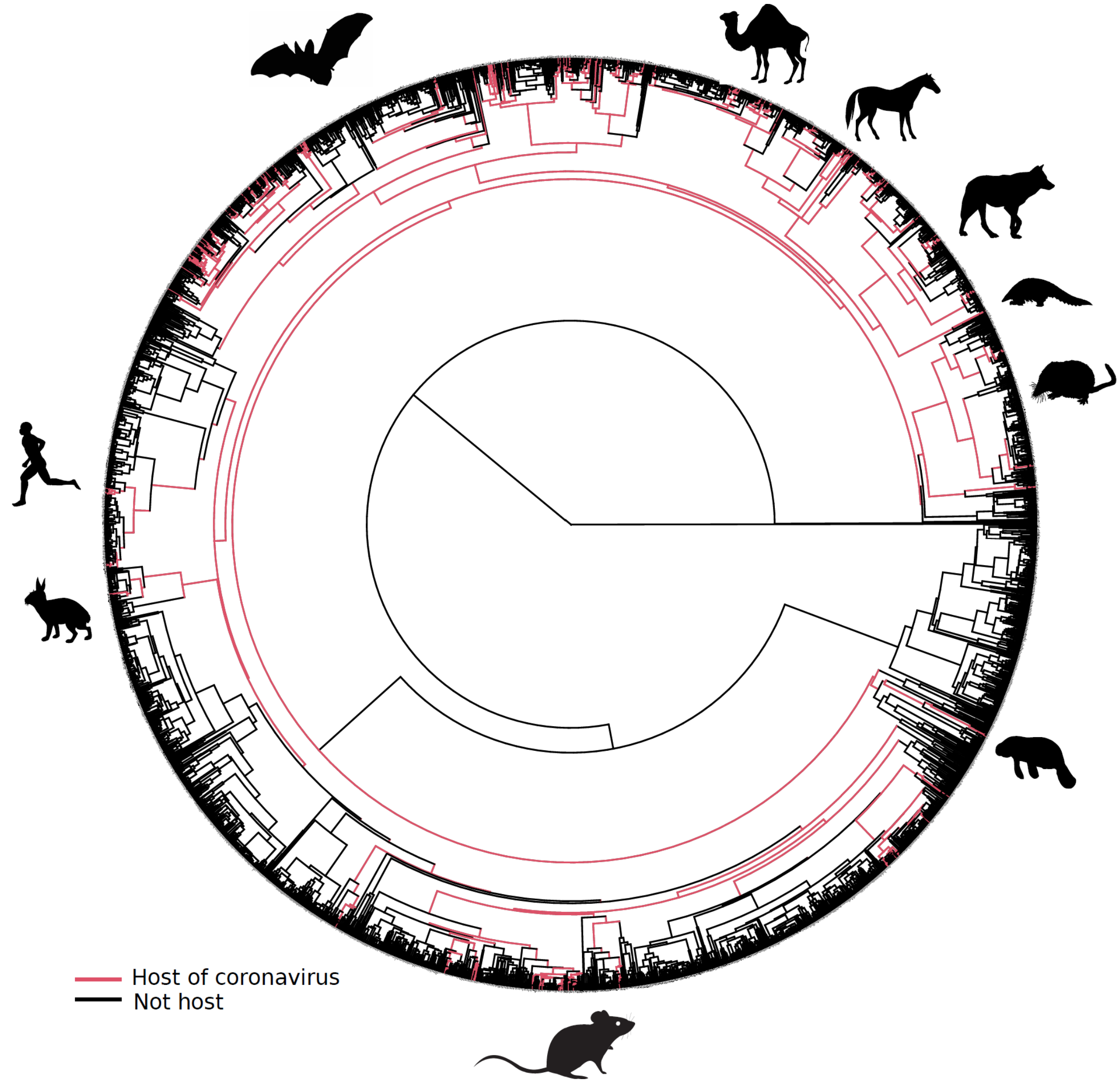


**Fig. S2. Mammalian hosts of coronaviruses are shown within the full mammalian tree.** Mammalian tree with tip branches painted in red according to the species that are hosts of coronaviruses. Ancestral branches directly linked to the path toward terminal hosts were painted as well. Mammal silhouettes taken from open-to-use sources in phylopic.org, detailed credits given in SI Appendix Table S6.

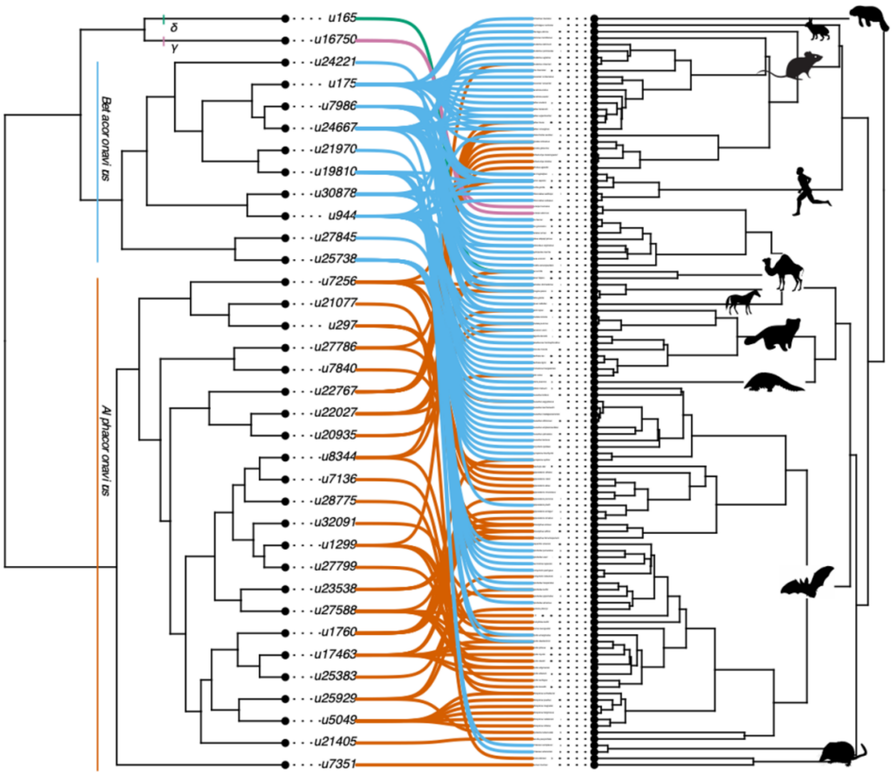


**Fig. S3. The association between coronaviruses and their mammalian hosts.** Coronaviruses tree on the left and mammalian tree on the right. The connections between coronaviruses and their hosts are shown with lines. Lines of different colors indicate different genera of coronaviruses (Betacoronaviruses in blue, Alphacoronaviruses in Orange). Mammal silhouettes taken from open-to-use sources in phylopic.org, detailed credits given in SI Appendix Table S6.

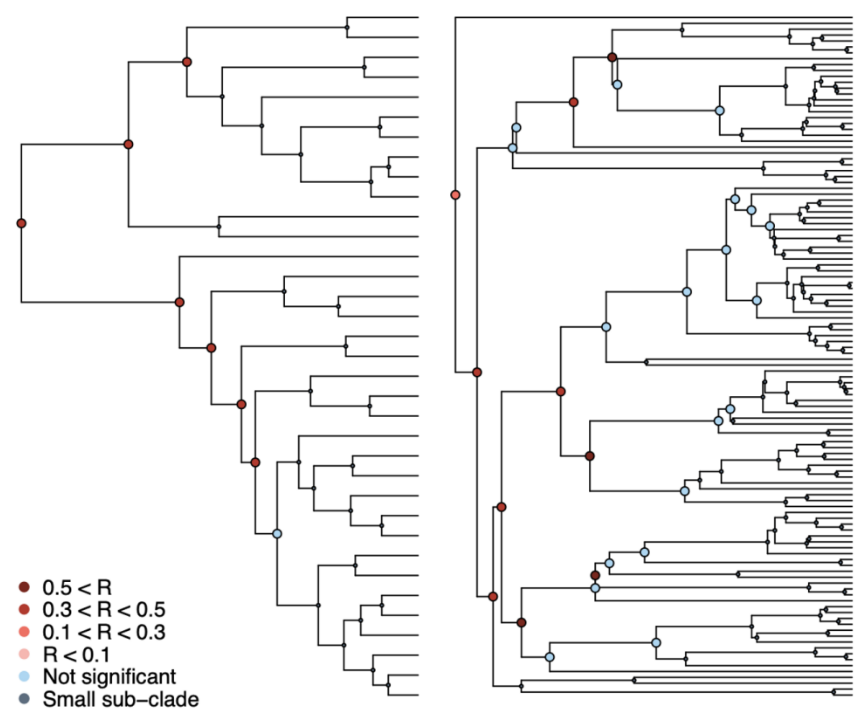


**Fig. S4. Phylogenetic signal in the association between coronaviruses and their mammalian hosts.** Coronaviruses tree on the left and mammalian tree on the right. Subclades tested for phylogenetic signal in the association matrix using Mantel tests are shown (see main text).

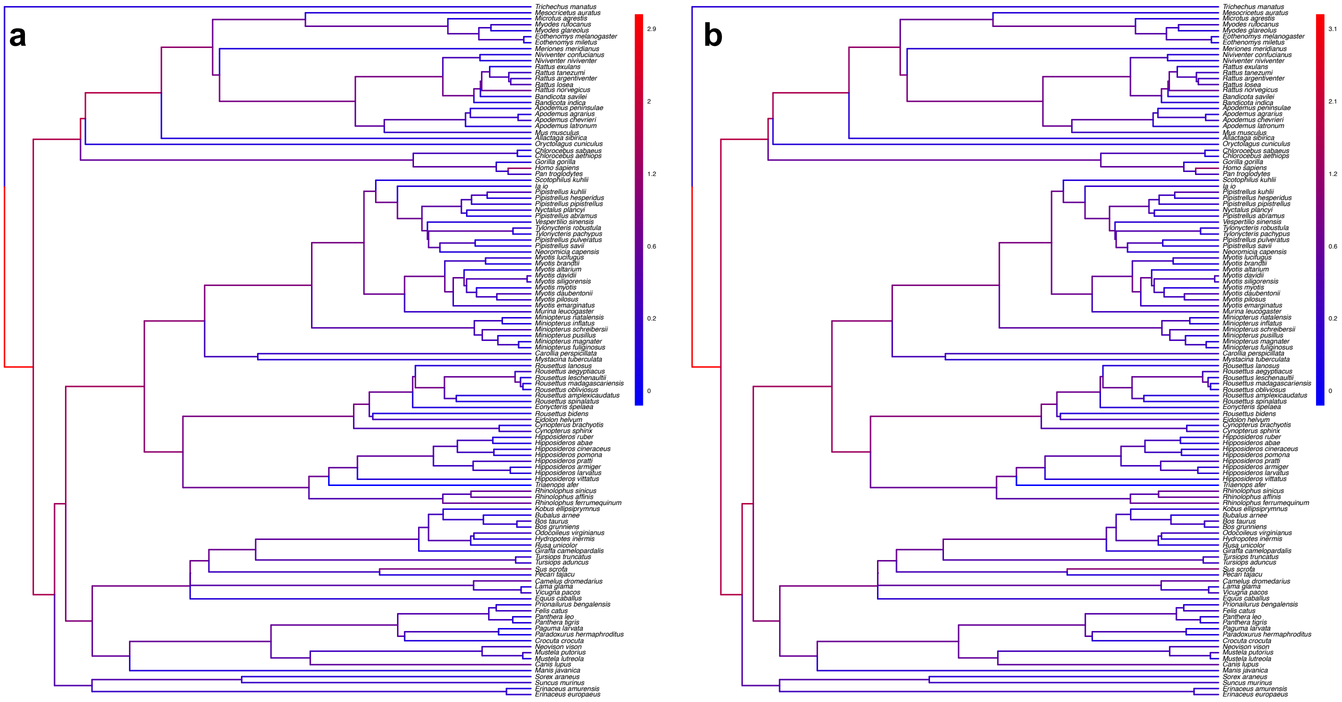


**Fig. S5. The origination of coronaviruses in mammals is not estimated among bats anymore when shuffling the dataset.**

Phylogenetic trees of the mammals with branches colored as the percentages of ALE reconciliations which inferred this branch as the origination of coronaviruses in mammals when the dataset is randomly shuffled (**a**) or shuffled by conserving mammal biogeography (by only shuffling species belonging to the same biogeographic realm; (**b**). Red branches are likely originations, whereas blue branches are unlikely.

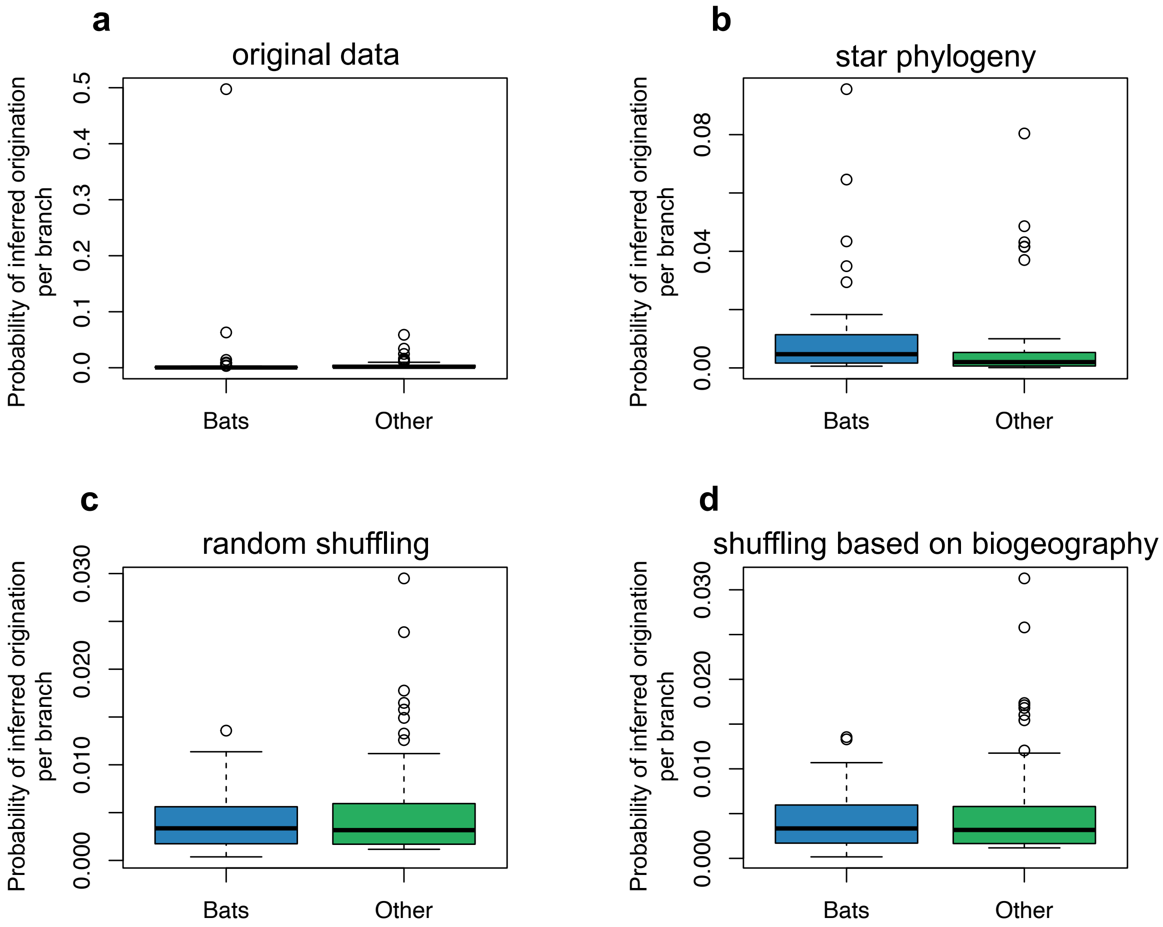


**Fig. S6. The origination of coronaviruses in mammals is estimated among bats.**
The boxplots indicated the probability of inferred origination per branch based on ALE reconciliations for bats lineages or non-bat lineages.
ALE was either run on: (**a**) the original dataset with the mammal phylogeny, (**b**) a star phylogeny instead of the mammal phylogeny, (**c**) on randomly-shuffled datasets, or (**d**) on datasets shuffled based on mammal biogeography (i.e. by only shuffling species belonging the same biogeographic realm).

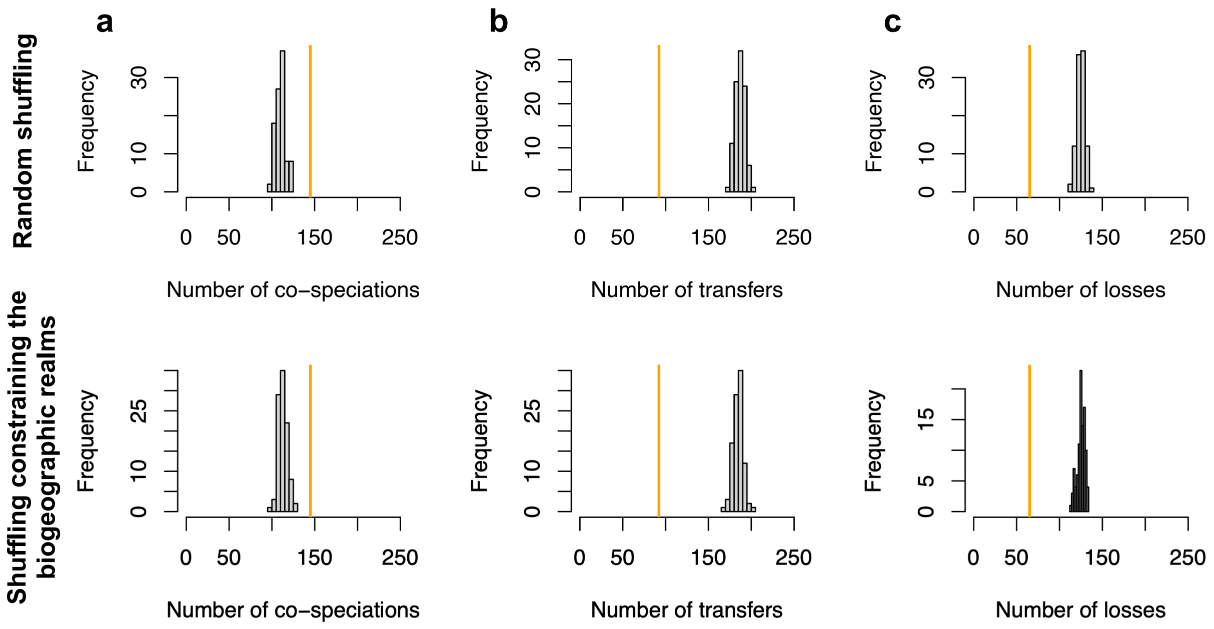


**Fig. S7. ALE inferred significant reconciliations.**
The number of cospeciation events (a), number of host switches (b), or number of losses (c) estimated on the original dataset (in orange) are significantly different from the numbers of events inferred when randomly shuffling the dataset (grey histograms; top row) or when shuffling by conserving mammal biogeography (by only shuffling species belonging the same biogeographic realm; bottom row).

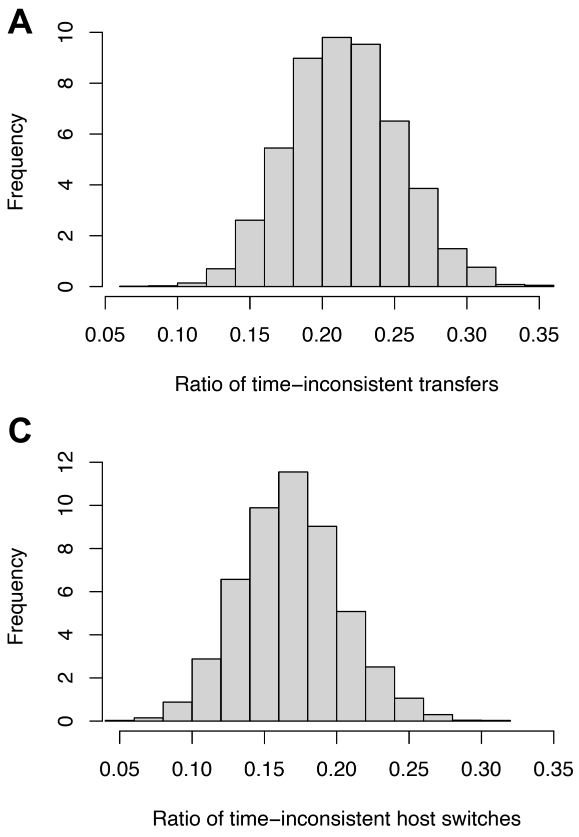


**Fig. S8. ALE inferred a large proportion of time-inconsistent host switches (a-b), which can be not be explained by the uncertainty in node age estimates (c-d)**

(a) Histogram of the percentage of time-inconsistent host switches in each reconciliation obtained with ALE on the original dataset with the consensus mammal phylogeny.

(b) Histogram of the time inconsistency (in Myr) of the inconsistent host switches.

(c) Histogram of the percentage of time-inconsistent host switches in each reconciliation obtained with ALE on the original dataset accounting for the 95% credible interval of the node age estimates. Although the mean number of time-inconsistent host switches decreased from 20% to 17% (meaning that 3% of the time-inconsistent host switches may be due to uncertainty in node age estimates), the reconciliations still contain frequent and large time-inconsistencies.

(d) Histogram of the time inconsistency (in Myr) of the inconsistent host switches accounting for the 95% credible interval of the node age estimates.

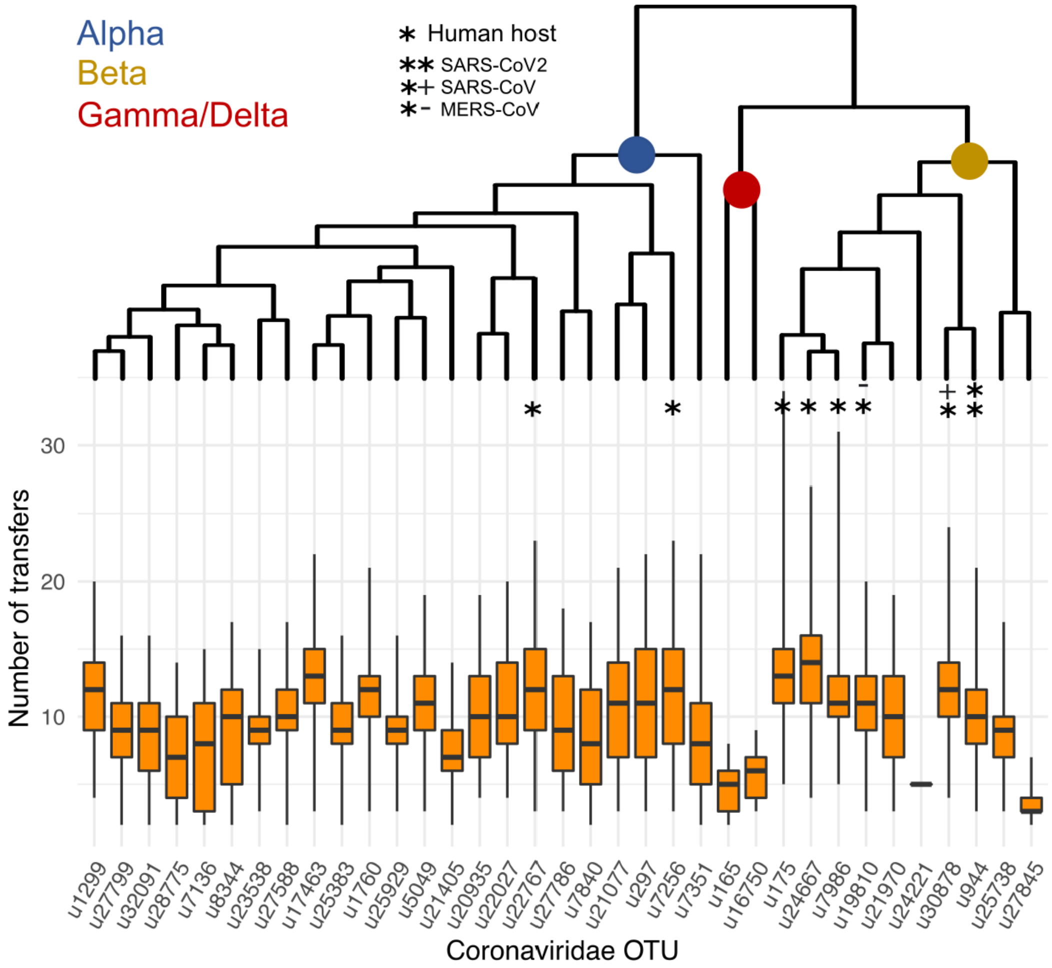


**Fig. S9. The frequency of host switches seems to vary according to the coronavirus lineages.**

On the top, we represented the phylogenetic tree of the coronavirus sOTUs reconstructed using BEAST2. For each extant sOTUs, we reported using boxplots, the total number of host switches that this OTU experienced since the coronavirus MRCA based on ALE reconciliations performed on a star phylogeny of mammals.

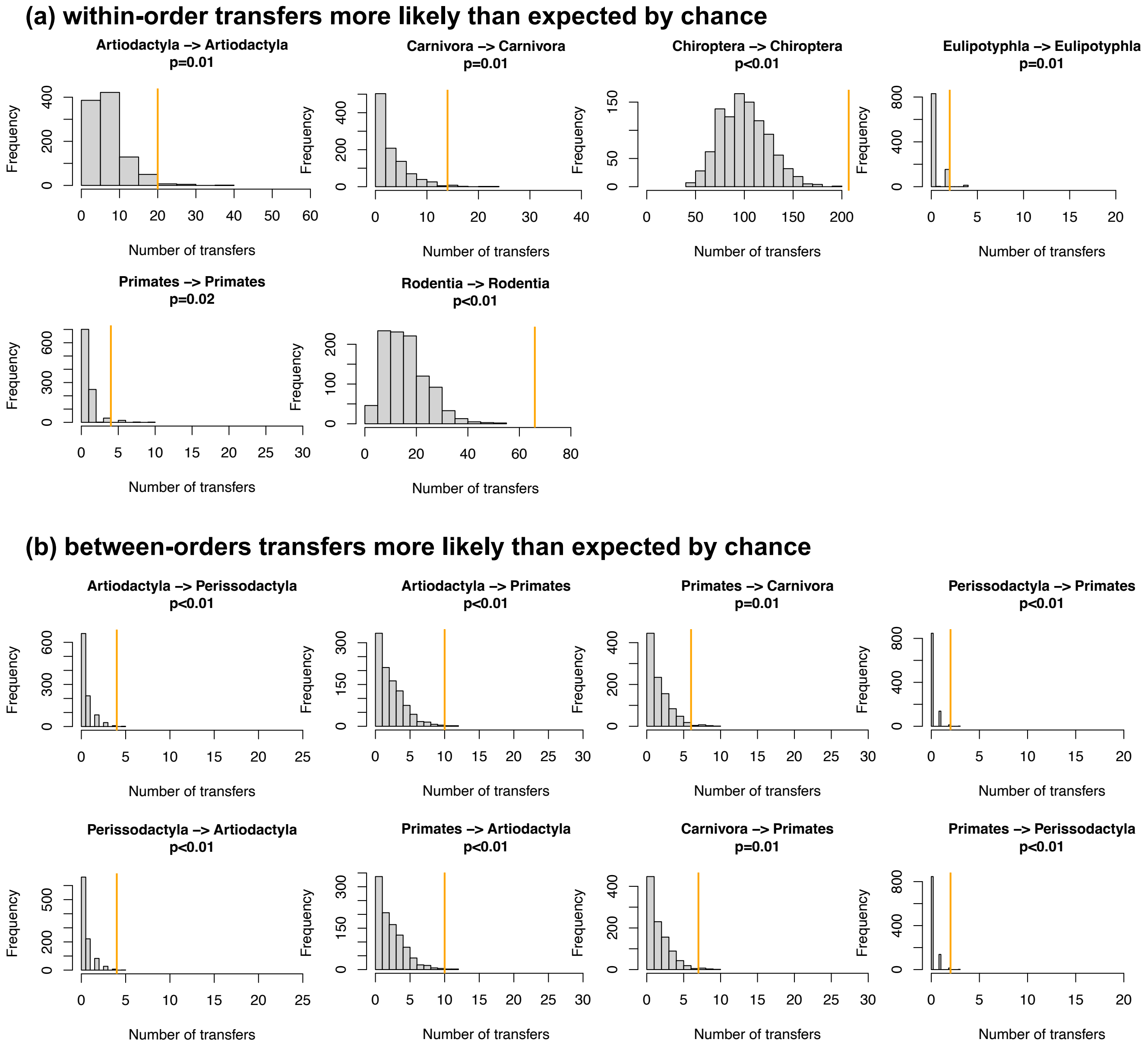


**Fig. S10. Evidence of preferential host switches in coronaviruses.**
(**a**) Numbers of **within**-order host switches estimated by ALE on the star mammal phylogeny (in orange) compared with the null expectations if host switches happen at random (grey histogram; obtained when randomizing the mammal species).
(**b**) For some clades, the numbers of **between**-order host switches estimated by ALE on the star mammal phylogeny (in orange) are higher than expected by chance.

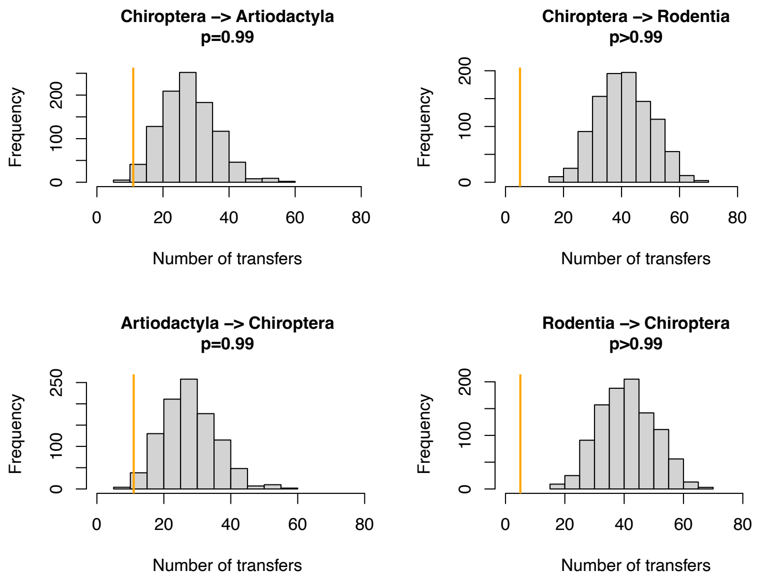


**Fig. S11. Host switches are less likely than expected by chance between bats (Chiroptera) and Artiodactyla or Rodentia.**
Numbers of **between**-order host switches estimated by ALE on the star mammal phylogeny (in orange) compared with the null expectations if host switches happen at random (grey histogram; obtained when randomizing the mammal species).

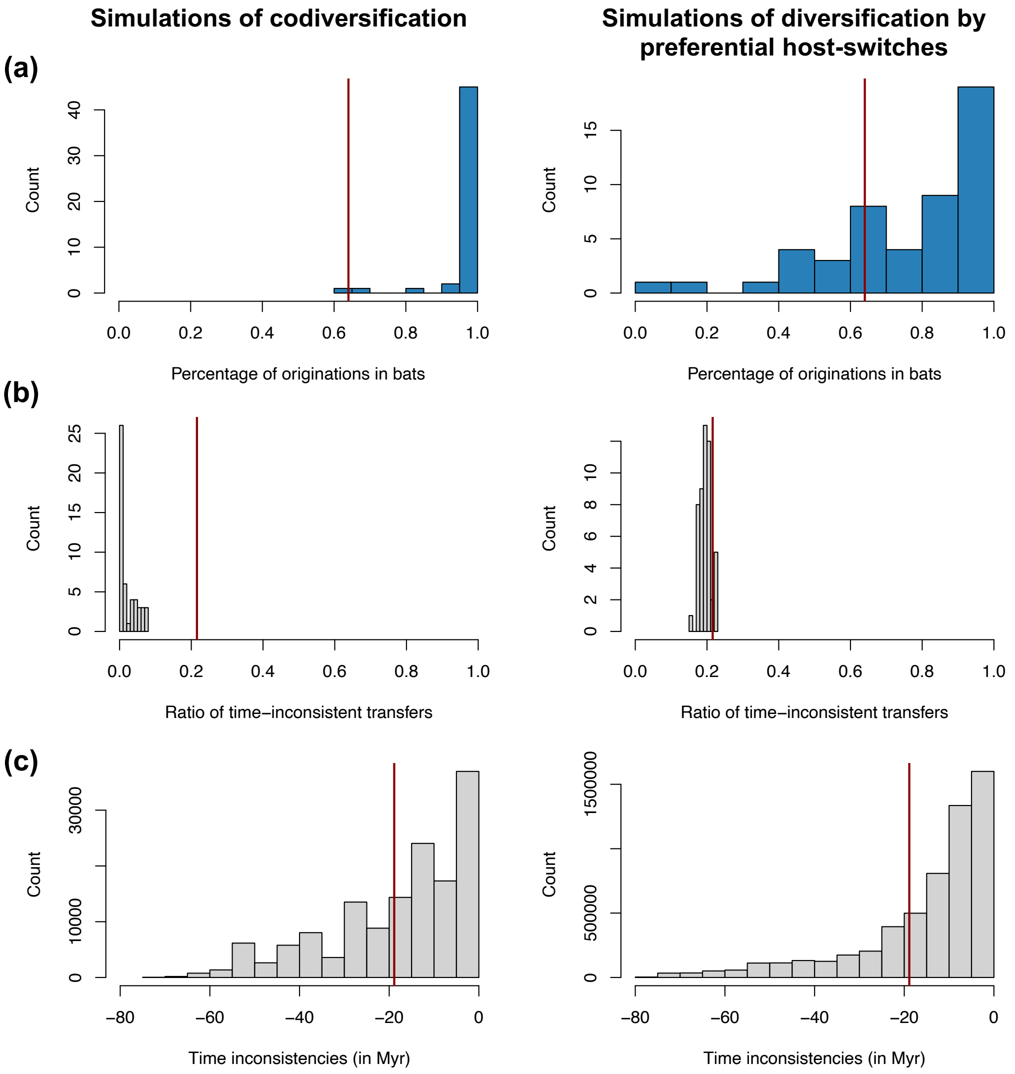


**Fig. S12. Validation of the interpretation of our results on the mammalian phylogeny using simulations of codiversification (left) or diversification by preferential host switches (right):** For each type of simulation – coronavirus-mammal codiversification (left) or coronavirus diversification by preferential host switches (right) –, we performed 50 independent simulations, ran ALE on the mammal phylogeny, and reported:

(a) The percentage of reconciliations inferring an origination within bats.

(b) The ratio of time-inconsistent host switches.

(c) Time inconsistencies (in Myr).

When simulating codiversification, ALE correctly infers an origination within bats, and few time-inconsistent host switches; when simulating preferential host switches, ALE correctly infers an origination within bats but with less certainty, and a significant fraction of time-inconsistent host switches.

For each plot, the vertical red line corresponds to the results obtained on the original data (empirical mammal-coronaviruses associations) using ALE on the mammal phylogenetic tree.

Results on the mammalian tree are consistent with a scenario of recent origination within bats and preferential host switches.

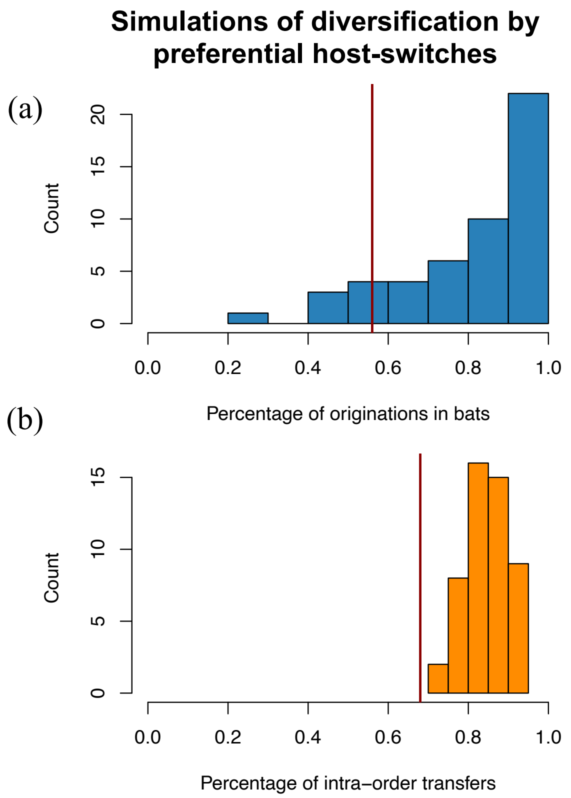


**Fig. S13. Validation of the interpretation of our results on the star phylogeny using simulations of diversification by preferential host switches:**

For each type of simulation – coronavirus-mammal codiversification (left) or coronavirus diversification by preferential host switches (right) –, we performed 50 independent simulations, ran ALE on a star phylogeny, and reported:

(a) The percentage of reconciliations happening within bats.

(b) The percentages of within-order host switches.

When simulating preferential host switches, ALE correctly infers a significant fraction of preferential host switches.

For each plot, the vertical red line corresponds to the results obtained on the original data (empirical mammal-coronaviruses associations) using ALE on a star phylogeny**.**

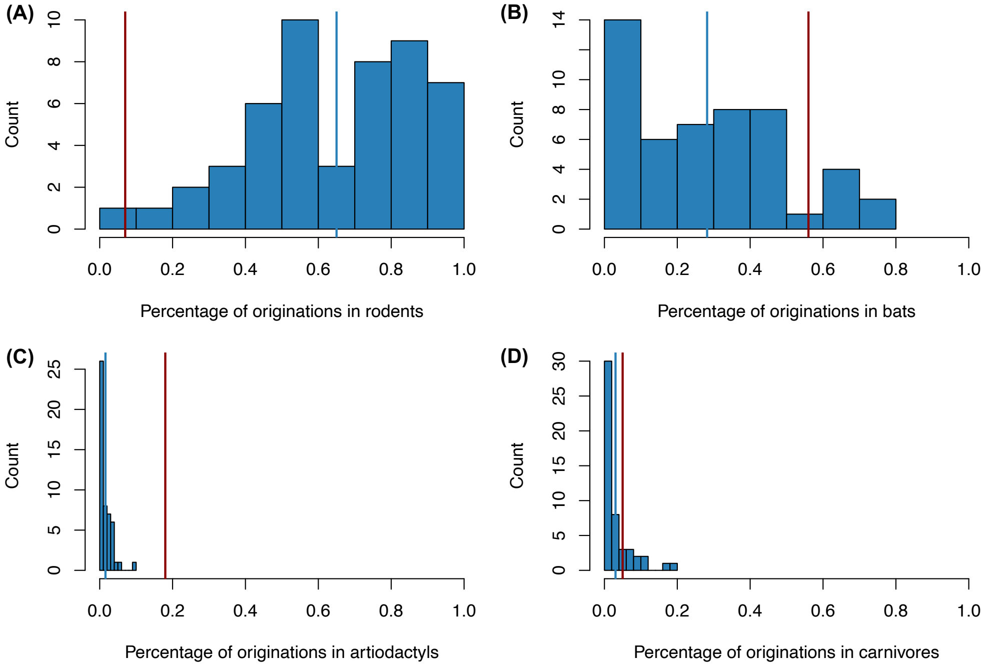


**Fig. S14. Simulating a scenario of origination in rodents followed by a diversification by preferential host switches with higher diversification of coronaviruses within bats did not generate a spurious origination in bats:**

We performed 50 independent simulations, ran ALE on a star phylogeny, and reported the percentage of originations within the main mammalian orders: rodents (A), bats (B), artiodactyls (C), and carnivores (D).

For each plot, the vertical red line corresponds to the results obtained on the original data (empirical mammal-coronaviruses associations) using ALE on a star phylogeny, while the vertical blue line corresponds to the mean of the simulations.

Originations were correctly inferred in rodents in the majority of the simulations (average percentage: 65% +/- s.d. 22%), and only in a minority of cases within bats (average percentage: 28% +/- s.d. 21%), artiodactyls (average percentage: 2% +/- s.d. 2%), or carnivores (average percentage: 3% +/- s.d. 4%).

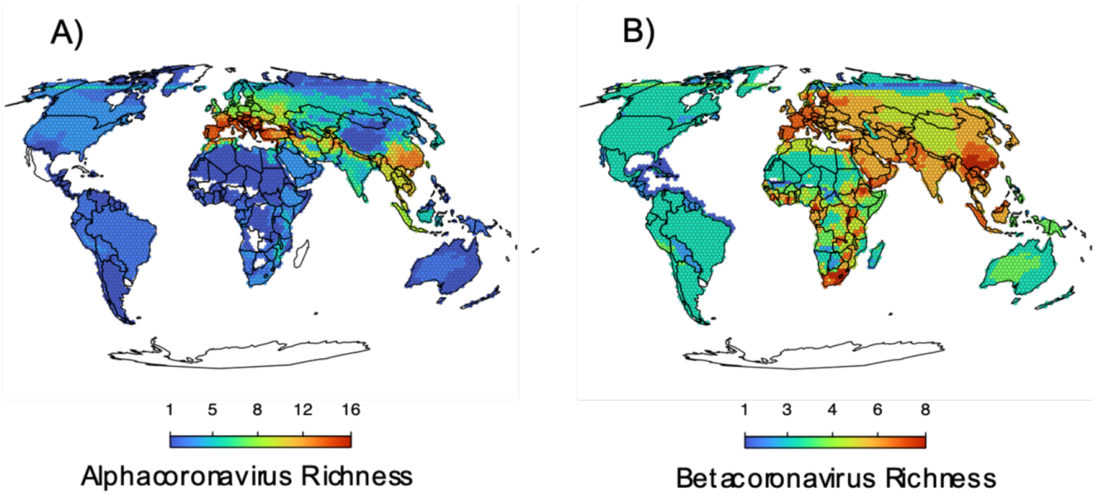


**Fig. S15: Maps of the diversity of alpha and betacoronaviruses.** In A) the richness of sOTUs of alphacoronaviruses and in B) the richness of sOTUs of betacoronaviruses; geographic range maps of coronaviruses were constructed after applying the host-filling method on the geographic range maps of mammalian hosts of coronaviruses.

**Table S1. Summary of the results obtained for the ALE reconciliations performed on "sliced mammalian phylogenetic trees", i.e.** trees where we only considered the X last Myr and merged nodes older than X Myr into polytomies in order to (i) test a scenario of more or less recent origination of coronaviruses and (ii) avoid back-in-time transfers toward nodes older than the origination time.
All reconciliations are significant (when compared to randomizations shuffling host species labels). Reconciliations with an estimated number of cospeciation events larger than the estimated number of transfer events are in bold.

| Sliced age of the mammalian phylogeny (in Myr ago) | Estimated numbers of cospeciations and transfers | Percentage of origination in bats | Percentage of time-inconsistent transfers | Average time inconsistency (in Myr) |
|---|---|---|---|---|
| 55 | 119<br>97 | 77% | 19% | -13 Myr |
| 50 | 113<br>98 | 85% | 18% | -11 Myr |
| 45 | 113<br>100 | 84% | 17% | -11 Myr |
| 40 | 110<br>102 | 83% | 17% | -10 Myr |
| 35 | 102<br>105 | 82% | 16% | -8 Myr |
| 30 | 100<br>106 | 83% | 15% | -7 Myr |
| 25 | 88<br>109 | 40% | 14% | -6 Myr |
| 20 | 87<br>110 | 42% | 15% | -5 Myr |
| 15 | 75<br>115 | 38% | 13% | -5 Myr |
| 10 | 45<br>127 | 32% | 8% | -4 Myr |
| 5 | 19<br>153 | 40% | 1% | -1.5 Myr |

**Table S2. Host switches between bats and other mammal orders are less likely than expected by chance:**

For each type of host switches within or between orders, we reported the inferred number of host switches using ALE (on the star phylogeny) and the expected number of host switches if host switches are equally likely between species (obtained by randomly shuffling the host species names). Because we don't have within-OTUs variations with the palmprint region, the directionality all the recent host switches is not identifiable (resulting in equal proportion in both directions at the mammalian order level).

Host switches involving bats (Chiroptera) and other mammal orders are indicated in bold.

| Type of host switch | Number of inferred host switches | Number of expected host switches if by chance |
|---|---|---|
| Chiroptera --> Chiroptera | 206 | 101 |
| Rodentia --> Rodentia | 66 | 16 |
| Artiodactyla --> Artiodactyla | 20 | 7 |
| Carnivora --> Carnivora | 14 | 4 |
| **Artiodactyla --> Chiroptera** | **11** | **28** |
| **Chiroptera --> Artiodactyla** | **11** | **28** |
| Artiodactyla --> Primates | 10 | 3 |
| Primates --> Artiodactyla | 10 | 3 |
| Primates --> Rodentia | 8 | 4 |
| Rodentia --> Primates | 8 | 4 |
| Artiodactyla --> Carnivora | 7 | 6 |
| Carnivora --> Primates | 7 | 2 |
| Primates --> Carnivora | 6 | 2 |
| Artiodactyla --> Rodentia | 5 | 11 |
| Carnivora --> Artiodactyla | 5 | 5 |
| **Chiroptera --> Rodentia** | **5** | **41** |
| **Primates --> Chiroptera** | **5** | **10** |
| Rodentia --> Artiodactyla | 5 | 11 |
| **Rodentia --> Chiroptera** | **5** | **41** |
| Artiodactyla --> Perissodactyla | 4 | infrequent |
| **Chiroptera --> Eulipotyphla** | **4** | **8** |
| **Chiroptera --> Primates** | **4** | **10** |
| **Eulipotyphla --> Chiroptera** | **4** | **7** |
| Perissodactyla --> Artiodactyla | 4 | infrequent |
| Primates --> Primates | 4 | 1 |
| Carnivora --> Rodentia | 3 | 8 |
| Rodentia --> Carnivora | 3 | 8 |
| Eulipotyphla --> Eulipotyphla | 2 | infrequent |
| Perissodactyla --> Primates | 2 | infrequent |
| Primates --> Perissodactyla | 2 | infrequent |

| | | |
|---|---|---|
| Carnivora --> Perissodactyla | 1 | infrequent |
| Eulipotyphla --> Primates | 1 | 1 |
| Perissodactyla --> Carnivora | 1 | infrequent |
| Perissodactyla --> Rodentia | 1 | 1 |
| Primates --> Eulipotyphla | 1 | 1 |
| Rodentia --> Perissodactyla | 1 | 1 |
| **Carnivora --> Chiroptera** | **infrequent** | **20** |
| **Chiroptera --> Carnivora** | **infrequent** | **20** |
| Eulipotyphla --> Rodentia | infrequent | 3 |
| Rodentia --> Eulipotyphla | infrequent | 3 |
| Artiodactyla --> Eulipotyphla | infrequent | 2 |
| Eulipotyphla --> Artiodactyla | infrequent | 2 |
| **Sirenia --> Chiroptera** | **infrequent** | **2** |
| **Chiroptera --> Sirenia** | **infrequent** | **2** |
| **Chiroptera --> Pholidota** | **infrequent** | **2** |
| **Pholidota --> Chiroptera** | **infrequent** | **2** |
| **Chiroptera --> Perissodactyla** | **infrequent** | **2** |
| **Perissodactyla --> Chiroptera** | **infrequent** | **2** |
| **Lagomorpha --> Chiroptera** | **infrequent** | **2** |
| **Chiroptera --> Lagomorpha** | **infrequent** | **2** |
| Eulipotyphla --> Carnivora | infrequent | 2 |
| Carnivora --> Eulipotyphla | infrequent | 2 |
| Pholidota --> Rodentia | infrequent | 1 |
| Rodentia --> Pholidota | infrequent | 1 |
| Rodentia --> Sirenia | infrequent | 1 |
| Sirenia --> Rodentia | infrequent | 1 |
| Rodentia --> Lagomorpha | infrequent | 1 |
| Lagomorpha --> Rodentia | infrequent | 1 |
| Pholidota --> Artiodactyla | infrequent | 1 |
| Artiodactyla --> Pholidota | infrequent | 1 |
| Artiodactyla --> Sirenia | infrequent | 1 |
| Sirenia --> Artiodactyla | infrequent | 1 |
| Artiodactyla --> Lagomorpha | infrequent | 1 |
| Lagomorpha --> Artiodactyla | infrequent | 1 |

**Table S3. Frequency of host switches inferred from bats to other mammal species, including humans.**

For each mammal species, we computed the average number per reconciliation of host switches from bats to this mammal species. We only reported here the species presenting >10% of chance to experience at least one host switch from bats.

| Mammal species | Average number of host switches per reconciliation from bats |
|---|---|
| *Homo sapiens* | 1.9 |
| *Rattus norvegicus* | 1.7288 |
| *Camelus dromedarius* | 1.4098 |
| *Sus scrofa* | 0.9996 |
| *Sorex araneus* | 0.8626 |
| *Vicugna pacos* | 0.7008 |
| *Lama glama* | 0.6912 |
| *Erinaceus amurensis* | 0.6258 |
| *Erinaceus europaeus* | 0.542 |
| *Suncus murinus* | 0.4996 |
| *Tursiops truncatus* | 0.168 |
| *Tursiops aduncus* | 0.1632 |
| *Canis lupus* | 0.1618 |
| *Mustela putorius* | 0.105 |

**Table S4. Frequency of host switches inferred from any mammal species towards humans.** We computed the average number per reconciliation of host switches from each mammal species to humans. We only reported here the species presenting >10% of chance to experience at least one host switch. Host switches from bats are highlighted in bold.

| Mammal species | Average number of host switches per reconciliation towards humans |
|---|---|
| *Camelus dromedarius* | 0.41 |
| *Mus musculus* | 0.31 |
| *Canis lupus* | 0.26 |
| *Sus scrofa* | 0.24 |
| *Suncus murinus* | 0.22 |
| *Paradoxurus hermaphroditus* | 0.21 |
| *Paguma larvata* | 0.21 |
| *Chlorocebus aethiops* | 0.21 |
| *Vicugna pacos* | 0.20 |
| ***Hipposideros vittatus*** | **0.17** |
| *Pan troglodytes* | 0.17 |
| *Bos taurus* | 0.17 |
| ***Rousettus aegyptiacus*** | **0.16** |
| ***Hipposideros abae*** | **0.16** |
| ***Hipposideros ruber*** | **0.15** |
| *Rattus losea* | 0.14 |
| *Rattus norvegicus* | 0.14 |
| *Myodes rufocanus* | 0.13 |
| *Equus caballus* | 0.13 |
| *Rattus tanezumi* | 0.12 |
| *Odocoileus virginianus* | 0.12 |
| *Rattus argentiventer* | 0.11 |
| *Neovison vison* | 0.10 |
| *Bos grunniens* | 0.10 |
| *Mustela putorius* | 0.10 |

**Table S5. Results are qualitatively similar when running ALE on sub-parts on the palmprint region.**

We reported here results obtained when running ALE on the star phylogeny on (i) the whole palmprint region (positions 1-150), (ii) the first part of the palmprint region (positions 1-75) or (iii) the last part of the palmprint region (positions 76-150).

| | **Whole palmprint region (positions 1-150)** | **First part of the palmprint region (positions 1-75)** | **Last part of the palmprint region (positions 76-150)** |
|---|---|---|---|
| Percentage of time-inconsistent host switches | 21% | 24% | 22% |
| Percentage of originations in bats | 56% | 46% | 64% |
| Percentage of within-order host switches | 68% | 70% | 68% |
| Percentage of host switches from bats to others (and *vice versa*) | 10% | 12% | 11% |

**Table S6.** Mammal silhouettes taken from open-to-use sources in phylopic.org, detailed credits for authors given below.

| **Mammal** | **Author** |
|---|---|
| Asian palm civet | Margot Michaud |
| Bat | Yan Wong |
| *Camelus dromedarius* | Steven Traver |
| *Equus caballus* | Jody Taylor |
| *Homo sapiens* | T. Michael Keesey |
| *Mus musculus* | Kamil S. Jaron |
| *Oryctolagus cuniculus* | Steven Traver |
| Pangolin | Steven Traver |
| *Sorex araneus* | Becky Barnes |
| Trichetus | Steven Traver |

**Table S7.** Summary of the different strategies used to evaluate the robustness of our findings.

| Potential bias or issues | Solution | Analyses |
|---|---|---|
| Over-sampling of humans and domesticated animals | Subsampling of the dataset to only 3 Genbank accessions per mammal species | PhyloBayes + ALE on each of the 50 subsampled datasets |
| Over-representation of bats in the dataset | Subsampling of the dataset (up to 10 species per mammalian order only) | PhyloBayes + ALE on each of the 50 subsampled datasets |
| The palmprint region of the RdRp region may be subject to recombination | Split of the palmprint region into two 150-amino acid subparts | PhyloBayes + ALE on each subpart |
| Coronavirus origination is spuriously inferred at the origin of Pteropodidae | Permutations of the host species in the mammal phylogeny (1) randomly or (2) by constraining by bioregion | PhyloBayes + ALE on each of the 100 randomized datasets |
| Origination cannot be inferred using a star phylogeny for the reconciliation | Simulations of scenarios of diversification per preferential host switches | PhyloBayes + ALE on each of the 50 simulated datasets |
| Observing 20% of time-inconsistent host switches can happen under a scenario of codiversification | Simulations of scenarios of codiversification | PhyloBayes + ALE on each of the 50 simulated datasets |
| Higher diversification rates of coronaviruses within bats can generate a spurious inference of an origination in bats | Simulations of scenarios of origination in rodents and diversification per preferential host switches with higher diversification rates of coronaviruses within bats | PhyloBayes + ALE on each of the 50 simulated datasets |

**Dataset S1 (separate file).** Association matrix between coronaviruses and their mammalian hosts.
